# The CLASSY family controls tissue-specific DNA methylation patterns in Arabidopsis

**DOI:** 10.1101/2021.01.23.427869

**Authors:** Ming Zhou, Ceyda Coruh, Guanghui Xu, Clara Bourbousse, Alice Lambolez, Julie A. Law

## Abstract

DNA methylation shapes the epigenetic landscape of the genome, plays critical roles in regulating gene expression, and ensures transposon silencing. As evidenced by the numerous defects associated with aberrant DNA methylation landscapes, establishing proper tissue-specific methylation patterns is critical. Yet, how such differences arise remains a largely open question in both plants and animals. Here we demonstrate that CLASSY1-4 (CLSY1-4), four locus-specific regulators of DNA methylation that are differentially expressed during plant development, play major roles in controlling tissue-specific DNA methylation patterns. Depending on the tissue, the genetic requirements for specific CLSYs differ significantly and, on a global scale, certain *clsy* mutants are sufficient to largely shift the epigenetic landscape between tissues. Together, these findings not only reveal substantial epigenetic diversity between tissues, but assign these changes to specific CLSY proteins, revealing how locus-specific targeting combined with tissue-specific expression enables the CLSYs to generate epigenetic diversity during plant development.

## Introduction

Global DNA methylation patterns reflect a balance between pathways controlling the establishment, maintenance and removal of this otherwise stable chromatin modification. Modulation within these pathways thus affords the opportunity to generate the unique DNA methylation patterns observed between somatic or reproductive tissues in plants^1,2^ and mammals^3-5^. For example, during plant reproduction, both male and female tissues (the vegetative nucleus and the central cell, respectively) are hypomethylated due to reduced maintenance and increased demethylation activities^2^. In somatic plant tissues, the differences in DNA methylation patterns are less well characterized. Nonetheless, key roles for the RNA-directed DNA methylation (RdDM) pathway, which controls the establishment of DNA methylation, are already emerging. Several recent studies in *Arabidopsis thaliana* (Arabidopsis)^6-9^, rice^10^, and soybean^11^ that compare the epigenomes of different somatic tissues and/or cell types all implicate the RdDM pathway in establishing cell type-specific DNA methylation patterns to differing extents. Furthermore, many of these studies suggest that there may be more differences in methylation patterns between plant tissues than previously envisioned^1,12^. However, in each of these cases, how the RdDM pathway is regulated to generate the observed tissue-or cell type-specific DNA methylation patterns remains unclear.

Much of our understanding of how DNA methylation is established and maintained stems from the characterization of proteins discovered using genetic and biochemical approaches. These studies have revealed multiple interconnected methylation and demethylation pathways that now provide a framework for understanding how different DNA methylation patterns are generated. Briefly, the RdDM pathway utilizes the sequence information encoded within two types of non-coding RNAs (24nt-siRNAs generated by RNA POLYMERASE-IV (Pol-IV) and long intergenic transcripts generated by Pol-V) to establish DNA methylation in all contexts (CG, CHG, and CHH; H=A, T or C) at cognate sequences throughout the genome^13,14^. Once established, DNA methylation in each context is maintained via largely distinct pathways involving self-reinforcing loops between DNA and/or histone methylation readers and writers^15,16^: CG methylation is maintained by METHYLTRANSFERASE 1 (MET1) in connection with the VARIATION IN METHYLATION (VIM1-3) family of DNA-methylation readers^17-19^. CHG methylation is maintained by CHROMOMETHYLTRANSFERASE 3 (CMT3) in connection with three SU(VAR)3-9 HOMOLOG proteins (SUVH4-6) that read CHG methylation and write H3K9 methylation^17,20-24^. Finally, CHH methylation at short, euchromatic transposons is maintained primarily by DOMAINS REARRANGED METHYLTRANSFERASE 2 (DRM2) in connection with SAWADEE HOMEODOMAIN HOMOLOG 1 (SHH1), an H3K9me reader^25,26^, while CHH methylation at long, pericentromeric transposons is primarily maintained by CMT2 in connection with SUVH4-6^27,28^. All contexts of DNA methylation are also subject to demethylation by methyl-cytosine glycosylases^29^. Together, these methylation and demethylation pathways constitute a dynamic system to control DNA methylation patterns.

Given the central role of 24nt-siRNAs in controlling where methylation is deposited, understanding how Pol-IV is regulated has been of keen interest and has already provided insights into how methylation patterns are modulated^16,30,31^. Initial investigation into the composition of the Pol-IV complex led to the identification of several co-purifying accessory factors^32,33^, including four related putative chromatin remodeling factors, CLASSY (CLSY) 1-4^32^. Recently, we demonstrated that the CLSYs act as locus-specific regulators of the RdDM pathway by facilitating Pol-IV recruitment to distinct genomic targets in connection with different chromatin modifications^34^. CLSY1 and CLSY2 are required for the association of SHH1 with the Pol-IV complex, linking Pol-IV targeting by these CLSYs to H3K9me2, whereas targeting by CLSY3 and CLSY4 relies on CG methylation, but not H3K9me2^34^. Here we describe tissue-specific roles for the four CLSY proteins. First, we show that tissues with different *CLSY* expression levels show distinct DNA methylation patterns and attribute these differences to the RdDM pathway. Next, we demonstrate that, depending on the tissue, different combinations of CLSYs, or even individual CLSY family members, control global 24nt-siRNA and DNA methylation patterns. For example, CLSY1 is required for the production of the vast majority of 24nt-siRNAs in leaf tissue, while CLSY3 is required for the production of high levels of 24nt-siRNAs at a few hundred loci that dominate the small RNA landscape in ovules. Finally, we found that while *clsy3,4*-dependent loci show more variance across tissues than *clsy1,2*-dependent loci, either of these double mutants is sufficient to shift the 24nt-siRNA landscape from one tissue to nearly the wild-type landscape of another. Together, these findings establish the CLSYs as major, tissue-specific regulators of plant epigenomes, providing mechanistic insight into how diverse DNA methylation patterns are generated during development.

## Results

### The *CLSYs* are unique amongst DNA methylation factors in their degree of differential expression during development

As a first step in determining whether the CLSYs play a role in shaping tissue-specific patterns of DNA methylation, their expression patterns were assessed during Arabidopsis development. Based on previously published data available in the ePlant database^35^, (**Fig. S1** and **Fig. 1A**), four tissues were selected for further characterization using GUS reporters driven by the *CLSY* promoters (**Fig. 1B**) and mRNA sequencing (mRNA-seq) (**Table S1** and **Fig. 1C**). Flower buds (Fl; stage 12 and younger), which contain tissues expressing all four *CLSY* genes; ovules (Ov; mature, but unfertilized), which are enriched for *CLSY3* and to a lesser extent *CLSY4*; 1^st^ and 2^nd^ true leaves (Lv; 25 day old plants) and rosettes (Rs; 15 day old, no roots), which are enriched for *CLSY1* (**Fig. 1**). Beyond showing the tissue-specific expression patterns of the *CLSYs*, further analysis of the mRNA-seq data revealed that these genes show the most diverse expression patterns when compared to the rest of the RdDM machinery, the main genes required for the maintenance of DNA methylation in the different sequence contexts, or the four demethylases **(Fig. 1C)**. These findings demonstrate that the CLSYs are excellent candidates for regulating tissue-specific DNA methylation patterns during plant development.

**Figure 1.**
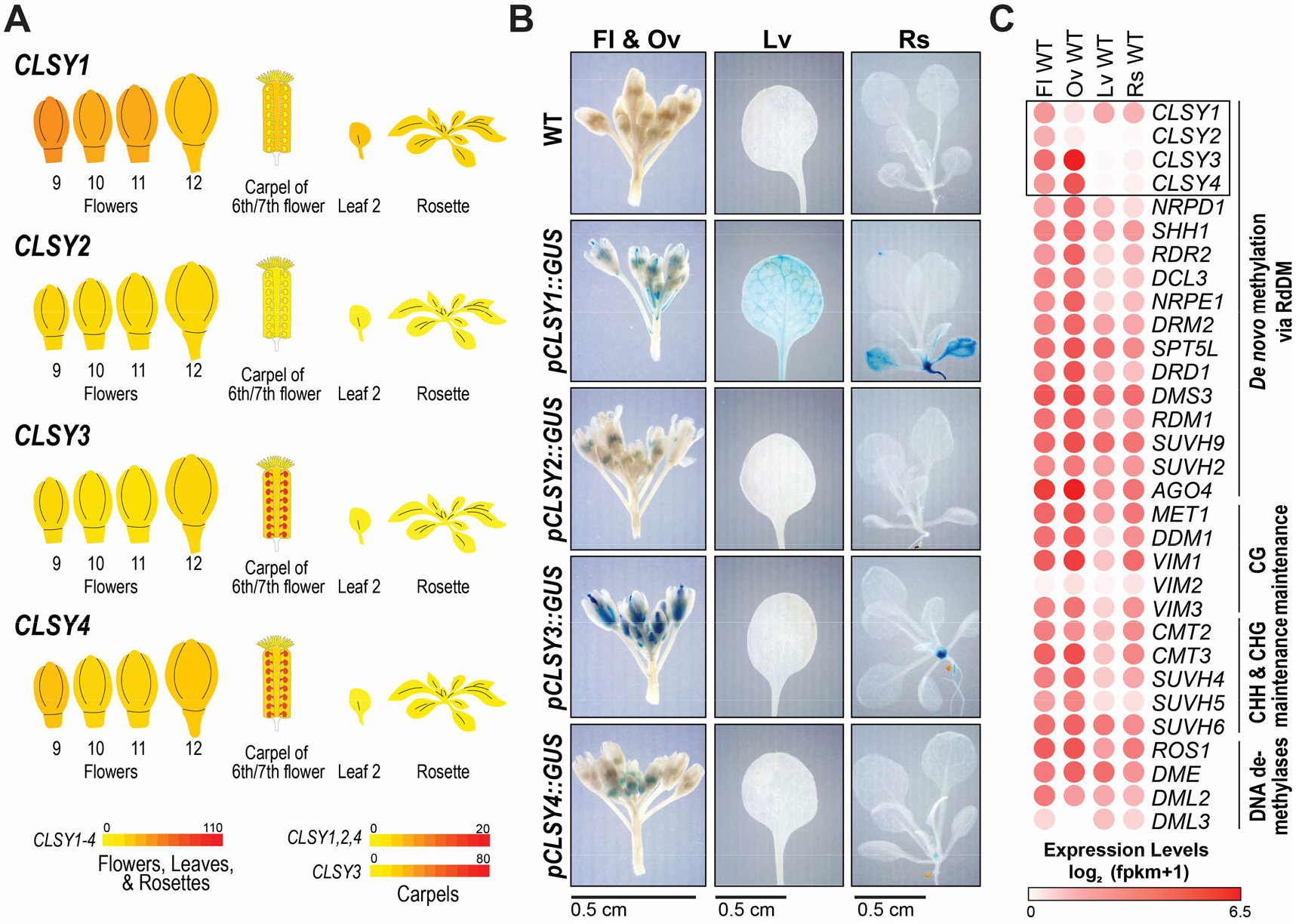
Tissue specific expression patterns of the *CLSY* genes.

(**A**) Relative expression levels of the four *CLSY* genes in select tissues from two ePlant expression viewers^68-70^ (see also **Fig. S1A, B**). Here, all tissues and organs are labeled as in the ePlant viewers^35^ and the cartoon images are not to scale. (**B**) Images showing GUS expression levels from wild-type (WT) plants or plants harboring GUS reporters driven by the promoters of the four CLSYs (*pCLSY1-4::GUS*) in the following tissues: flower buds (Fl; stage 12 and younger), unfertilized ovules (Ov), 1st and 2nd true leaves (Lv) from 25 day old plants, and rosettes (Rs; no roots) from 15 day old plants. For each set of images, a scale bar is shown below and for the true leaf samples, only a single representative leaf is shown. (**C**) Heatmap showing the expression levels of the main genes required for the establishment, maintenance, and removal of DNA methylation based on mRNA-seq data (**Table S1**) from same tissues described in panel **B**. The *CLSY* genes are surrounded by a black box.

### Tissues with different *CLSY* expression profiles have distinct epigenetic landscapes that are controlled by the RdDM pathway

Although DNA methylation patterns have been described for Arabidopsis flower, leaf, and rosette tissues individually, these patterns have not been compared to each other, and no DNA methylation data is available for ovules. Thus, to compare the epigenomes of these tissues, and assess the contribution of the RdDM pathway in shaping these landscapes, small RNA-seq (smRNA-seq) (**Table S2**) and MethylC sequencing (methylC-seq) (**Table S3**) data sets were generated in parallel from both wild-type samples and RdDM pathway mutants (*e*.*g. nprd1*; hereafter named *pol-iv*).

Initial comparisons of 24nt-siRNA levels on a chromosomal scale revealed distinct profiles amongst the four tissues and confirmed a strong dependency on *pol-iv* across all tissues (**Fig. 2A** and **Fig. S2A)**. Relative to flowers, the patterns in leaf and rosette tissues differ most in the chromosome arms and edges of the pericentromeric heterochromatin, where 24nt-siRNA levels are lower (**Fig. 2A** and **Fig. S2A**). However, the most striking difference was observed in ovules, where 24nt-siRNA levels are globally low with the exception of a small number of high expressing regions. For CHH methylation, which is known to be controlled by both the RdDM and CMT2 pathways^27,28^, the differences between flower, leaf, and rosette tissues in both the wild-type and *pol-iv* mutants are less evident at this scale (**Fig. 2A** and **Fig. S2A**). However, for ovules, the CHH methylation profiles were strongly *pol-iv*-dependent and highly correlated with 24nt-siRNA levels (**Fig. 2A** and **Fig. S2A**), demonstrating a major role for the RdDM pathway in this tissue. Taken together, these data demonstrate that epigenetic features associated with the RdDM pathway differ between flower, ovule, leaf, and rosette tissues to a degree that is apparent even on a genome-wide scale.

**Figure 2.**
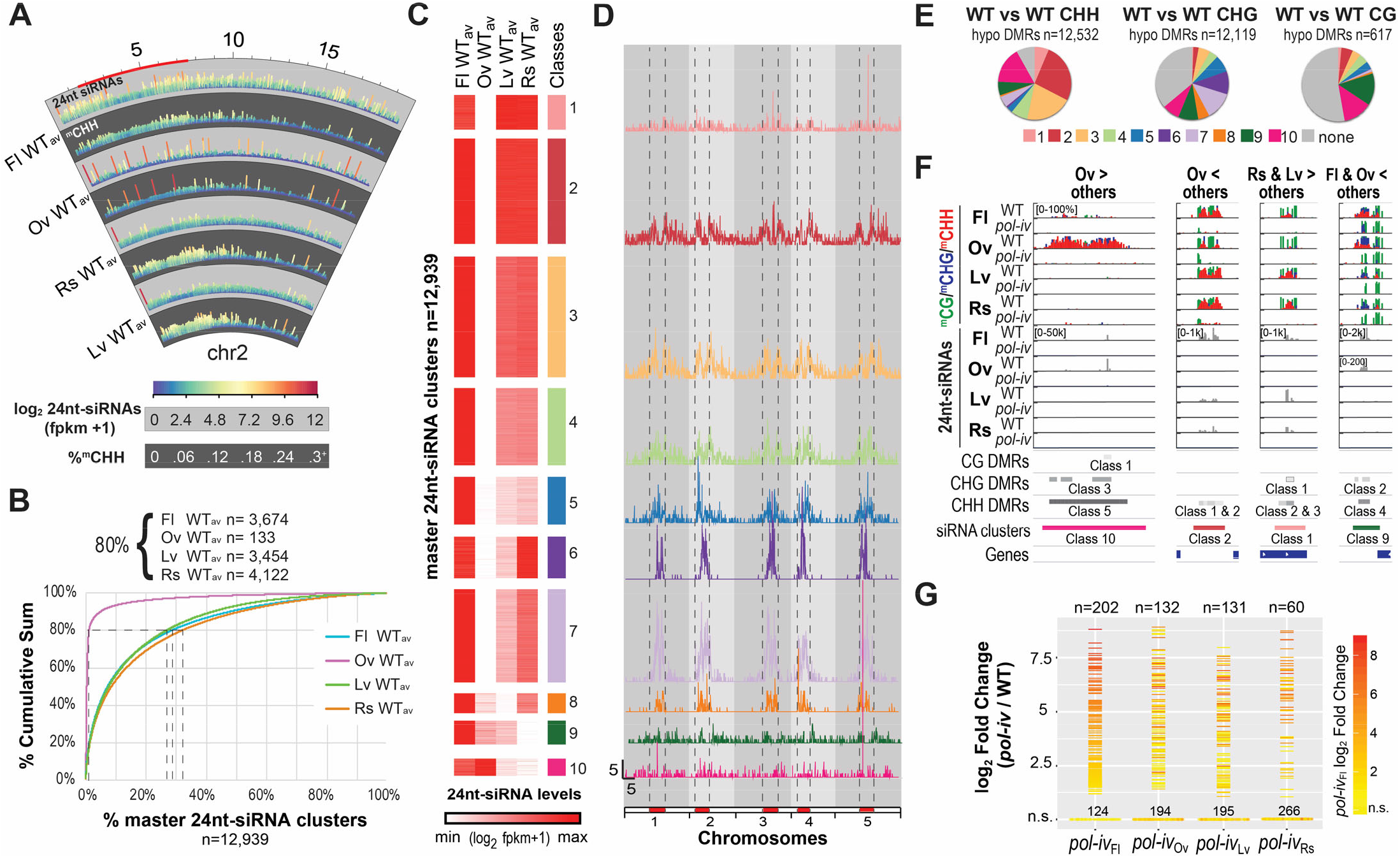
Tissues with different *CLSY* expression patterns have distinct epigenetic landscapes.

**A**) Circular genome view of chromosome 2 (chr2) showing the patterns of 24nt-siRNAs (light grey background) and CHH methylation (dark grey background) in 5kb bins based on the average (av) expression of three WT controls (WT_av_) from each tissue. The color scales for the data sets are as indicated and the tracks are labeled every 5Mbs, with the pericentromeric heterochromatin, as designated in Yelina *et al*.^66^, marked in red along the outer rim. For CHH methylation, between 1 and 3 bins, depending on the tissue, had values over 0.3 (0.3^+^), but were capped at this value to facilitate visualization on a genomic scale. (**B**) Cumulative sum plot based on the WT_av_ expression levels for each tissue, showing the percentage and number (n) of 24nt-siRNA clusters (x-axis) required to reach 80% of the fpkm-normalized 24nt-siRNA reads (y-axis) in each tissue. (**C**) Heatmap showing the designation of 10 classes of 24nt-siRNA clusters based on the WT_av_ expression levels for each tissue at the 12,939 master 24nt-siRNA clusters. (**D**) Distributions of 24nt-siRNA clusters from the classes described in **C** along the 5 chromosomes. The pericentromeric regions are marked in red and denoted by vertical dashed lines. Scale bars for x-axis (Mb) and y-axis (clusters/100kb bin) are indicated in the lower left corner of the Class 10 dataset. (**E**) Pie charts showing the proportions of hypo DMRs in each context that overlap with the 24nt-siRNA cluster classes colored as in **C**. (**F**) Screenshots showing the levels of 24nt-siRNAs and DNA methylation in all contexts at representative sites showing differential methylation between tissues. Track scales, indicated in brackets, apply to all lower tracks of the same tissue and DNA methylation in the CG, CHG, and CHH contexts are shown in green, blue, and red, respectively. DMRs, colored based on the classes in **Fig. S3A**, and 24nt-siRNA clusters, colored based on the classes in panel **C**, are indicated below. The greater than (>) and less than (<) symbols in the headers refer to relative DNA methylation levels. (**G**) Plot showing the expression levels [log_2_ FC (*pol-iv*/WT)] of the 326 *pol-iv*-upregulated transcripts represented as horizontal lines (log_2_ FC ≥ 1 and FDR ≤ 0.05) or dots (not significantly upregulated; n.s.) in the four tissues. The horizontal lines are colored based on the expression level in *pol-iv* from flower tissue (Fl).

To compare the profiles of these tissues in more detail, 24nt-siRNA clusters were first identified based on all three wild-type replicates from smRNA-seq experiments using ovule, leaf, and rosette samples, as was done previously for flowers^34^ (**Table S4**). The data from all four tissues were then merged to generate a list of “master 24nt-siRNA clusters” (n=12,939) representing all the clusters amongst the four tissues (**Table S5**). These 24nt-siRNA clusters were then grouped into 10 classes based on their expression levels and comparisons between the expression patterns and genomic distributions of each class revealed several tissue-specific trends (**Fig. 2C, D** and **Table S6**). First, in ovules, the levels of 24nt-siRNAs were low in all classes except 8-10, which show increasing expression levels, respectively (**Fig. 2C**). Indeed, the top 133 expressing clusters in ovules, which correspond well with the 24nt-siRNA peaks in the chromosome views (**Fig. 2A** and **Fig. S2A**), account for 80% of all the 24nt-siRNAs (**Fig. 2B** and **Table S5**). Given the similarity of these ovule loci to those described in rice^36^ and Brassica^37^, they are hereafter referred to by their initial designation as siren (small-interfering RNA in endosperm) loci^36^. Second, the expression profiles in Class 1, Class 2, and to a lesser extent Class 3, are similar in flower, rosette, and leaf tissues (**Fig. 2C**). These clusters, which correspond to nearly half of all 24nt-siRNA clusters (47%), appear to be similarly acted upon by Pol-IV across diverse tissues, which is consistent with their relative enrichment in the chromosome arms (**Fig. 2D**). Third, the expression levels in Classes 4-8 are much lower in leaf tissue compared to rosette and/or flower tissues (**Fig. 2C**), revealing that 24nt-siRNA production in pericentromeric heterochromatin (**Fig. 2D**) is less robust in leaves. Finally, while flowers show strong expression across all classes, leaf and rosette tissues, like ovules, show low expression in specific classes, namely Class 8 for leaf and Class 9 for rosette tissues (**Fig. 2C**). Together, these analyses reveal that tissue-specific differences in 24nt-siRNA patterns vary based on chromosomal locations and range from being essentially absent in a single tissue to being highly dynamic across tissues, suggesting a tunable means to control 24nt-siRNA patterns.

To determine the extent to which the observed tissue-specific differences in 24nt-siRNA levels contribute to altered DNA methylation patterns, differentially methylated regions (DMRs) between tissues were first determined in 100bp bins across the whole genome and then mapped onto the 24nt-siRNA landscape. Notably, this method required the identification of DMRs in all pairwise combinations of wild-type data sets (*e*.*g*. 3 wild-type ovule datasets versus 3 wild-type flower datasets = 9 pairwise combinations), and required 10%, 20%, or 40% changes for CHH, CHG and CG methylation, respectively (**Table S7**). Thus, rather than capturing the full range of epigenetic differences, it reveals the most robust tissue-specific differences in methylation between each pair of tissues. When combined, these analyses yielded a total of 12,532 CHH and 12,119 CHG DMRs that were hypo methylated between at least two tissues, but only 617 hypo CG DMRs (**Table S7**; “WTvsWT all tissues”). The majority of these DMRs overlap with 24nt-siRNA clusters (**Fig. 2E**) and, when clustered by their methylation levels, these DMRs show tissue-specific differences that are most apparent at loci regulated by *pol-iv* (**Fig. S3A** and **Table S8**). Thus, as exemplified by the regions shown in **Fig. 2F**, these analyses not only reveal that DNA methylation patterns differ significantly between these tissues at thousands of loci, especially for non-CG methylation, but they also demonstrate that these differences are largely attributable to the RdDM pathway.

Finally, to assess the role of the RdDM pathway in facilitating the silencing of genes and transposons in a tissue-specific manner, differentially expressed loci in *pol-iv* mutants were identified using RNA-seq data from flower^34^, ovule, rosette and leaf tissues and combined to generate a list of *pol-iv*-dependent loci amongst all tissues (**Tables S9** and **S10**). To visualize and compare the expression levels of these loci between tissues, a profile plot was generated where significantly upregulated loci (n=326; log_2_ Fold Change (FC) ≥ 1 and FDR ≤ 0.05; **Table S10**) are shown as bars, colored relative to the *pol-iv* levels in flowers, while those not passing these thresholds are shown at the bottom as dots (**Fig. 2G**). While there are some loci that are upregulated across all tissues (red/orange lines), there are others that are most strongly up in flowers, but are not significantly up in the other tissues (orange/red dots at the bottom of the plots) and *vice versa*. As was previously shown in flowers^34^, the vast majority of upregulated loci in all these tissues show proximal changes in DNA methylation and 24nt-siRNA levels (**Fig. S4A**), consistent with their mis-regulation being a direct effect of an altered epigenome. Together, these findings demonstrate that tissue-specific differences in gene silencing also rely on the RdDM pathway, highlighting major roles for this pathway in shaping the epigenomes of flower, ovule, leaf, and rosette tissues by controlling patterns of 24nt-siRNAs, DNA methylation, and gene expression.

### The CLSYs control 24nt-siRNA and DNA methylation patterns in both a locus- and tissue-specific manner

To determine what roles the CLSY proteins play in shaping the epigenomes of the aforementioned tissues, differentially expressed 24nt-siRNA clusters and DMRs were identified in *clsy* single, double and quadruple mutants relative to tissue-matched wild-type controls at the master set of 24nt-siRNA clusters (**Table S11**) or in 100bp bins (**Table S12**), respectively. The hypo CHH and, to a lesser extent, hypo CHG DMRs in each mutant and tissue were largely *pol-iv*-dependent (**Table S12**), similar to what we observed in the 24nt-siRNA clusters (**Fig. S2A**). Despite this dependence, many hypo CHH DMRs, especially in ovule and leaf tissues, did not overlap with reduced 24nt-siRNA clusters using our standard cutoff (log_2_ FC ≤-1; FDR ≤0.01). Investigation into this discrepancy revealed that 24nt-siRNAs were largely present at these DMRs, but their expression was very low in wild-type samples making further reduction in mutant backgrounds difficult to resolve statistically. Thus, to capture a larger proportion of regions controlled by the RdDM pathway in each tissue, 24nt-siRNA clusters that overlapped with *pol-iv*-dependent hypo CHH DMRs, and were reduced by at least 25% compared to wild-type controls (log_2_ FC < -0.415; **Fig. S5A**), were added to the lists of differentially expressed clusters for each tissue and genotype (**Table S11** and **Fig. 3A**). These regions were then used to assess both 24nt-siRNA and DNA methylation patterns across tissues and genotypes (**Fig. 3** and **Figs. S6-S9**).

**Figure 3.**
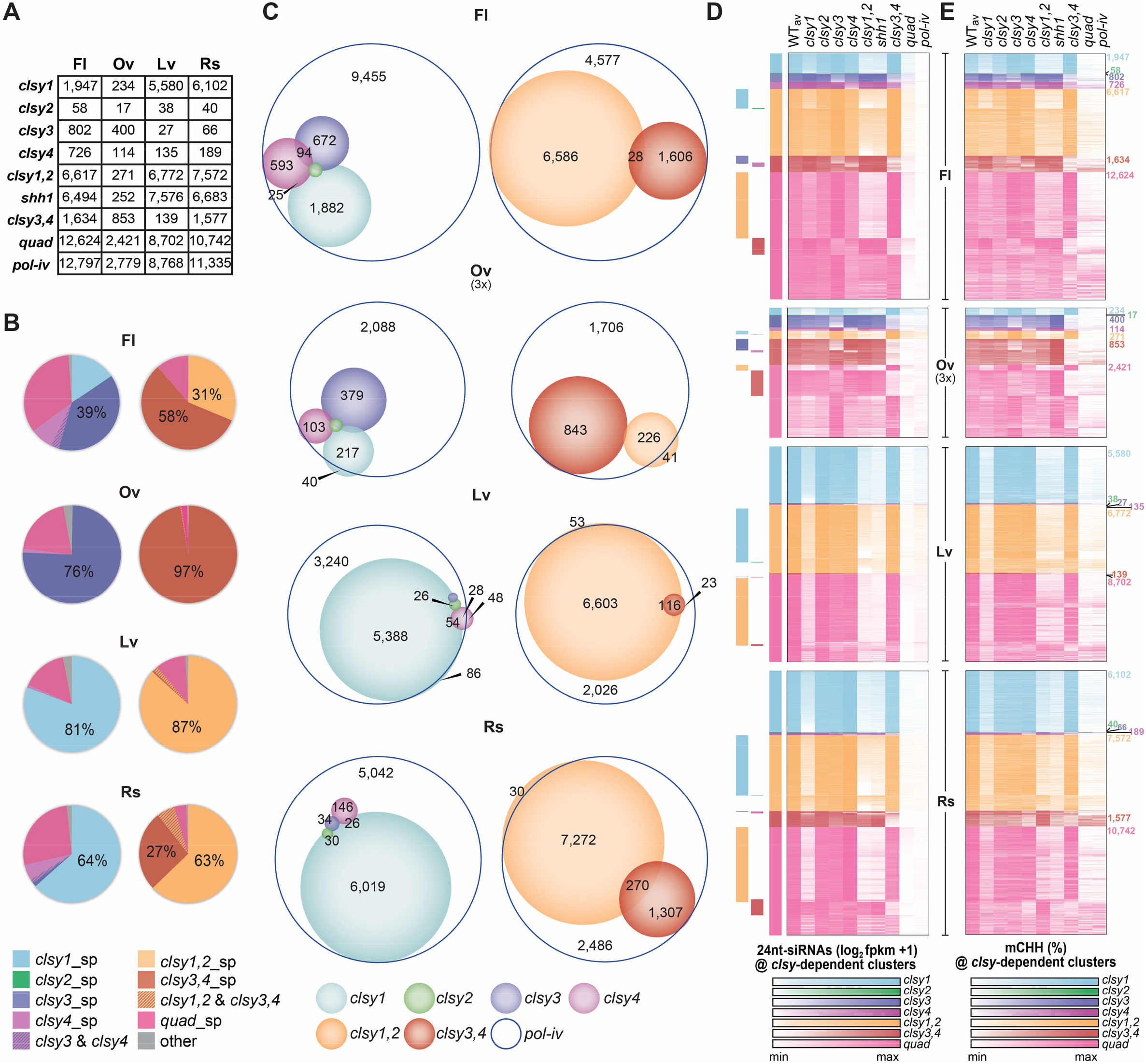
The CLSYs regulate 24nt-siRNA and DNA methylation patterns in a tissue-specific manner.

(A) Table showing the numbers of downregulated 24nt-siRNA clusters in the indicated tissues and genotypes. (**B**) Pie charts showing the proportions of *pol-iv*-dependent 24nt-siRNAs in each *clsy*-dependent category based on a WT control for each tissue and colored as indicated below. Here, “sp” indicates only 24nt-siRNAs clusters specifically reduced in the indicated genotype (*e*.*g*. not reduced in any other single mutant (left) or double mutants (right)) are included and “” indicates clusters that overlap between the indicated genotypes. (**C**) Scaled Venn diagrams showing the relationships between reduced 24nt-siRNA clusters in *pol-iv* and the *clsy* mutants colored as indicated below. For readability, only overlaps >20 are labeled. A small number of overlaps is not shown due to spatial constraints. Unscaled Venn diagrams showing all the overlaps is presented in **Fig. S5B**. (**D** and **E**) Heatmaps showing the 24nt-siRNA levels (**D**) or CHH methylation levels (**E**) for the WT_av_ or the indicated mutants at the reduced 24nt-siRNA clusters identified for each tissue. The heatmaps for each *clsy*-dependent category are colored as indicated below. For **D**, the heatmaps for each tissue are ranked (high to low) for each *clsy*-dependent category and the overlaps between categories are indicated by the colored bars on the left. For **E**, the heatmaps are ordered as in **D**. The number of clusters in each category are indicated on the right. For **C, D**, and **E** the data for ovules are scaled as 3x compared to the other tissues due to the reduced number of affected loci.

As a percentage of wild-type 24nt-siRNA abundance (**Fig. 3B**), reduced 24nt-siRNA clusters (**Fig. 3C**) or hypo CHH DMRs (**Fig. S6-9**), the four tissues show strikingly different degrees of dependence on the four CLSYs in a manner consistent with their observed expression patterns (**Fig. 1**). In terms of abundance, nearly all *pol-iv*-dependent 24nt-siRNAs in ovules reside in clusters reduced in *clsy3,4* mutants (**Fig. 3B**; 97%) while in leaves they reside in clusters reduced in *clsy1*,2 mutants (**Fig. 3B**; 87%). This is in contrast to rosette and flower tissues where there is a more even reliance on these CLSY pairs, although there is skew towards loci reduced in the *clsy1,2* mutant for rosette tissue and the *clsy3,4* mutant for flower tissue (**Fig. 3B**). Notably, when compared to flowers, most of the differences in these tissues stem from an increased proportion of 24nt-siRNAs in clusters reduced in *clsy1* mutants in rosette (64%) and leaf (81%) tissues or by those reduced in *clsy3* mutants in ovule (76%) tissue (**Fig. 3B**), highlighting the impact even a single CLSY protein can have in controlling 24nt-siRNA populations in a tissue-specific manner.

In terms of 24nt-siRNA clusters and hypo CHH DMRs, their patterns are similarly diverse between tissues and are largely in agreement with the 24nt-siRNA abundance data. For example, in rosette and leaf tissues the largest numbers of reduced 24nt-siRNA clusters and hypo CHH DMRs are observed in the *clsy1* mutant and relatively few sites were affected in the *clsy2, clsy3* or *clsy4* mutants (**Fig. 3C** and **Fig. S6-7**), reinforcing the dominance of CLSY1 in shaping the epigenomes of leaf and rosette tissues. Here, the difference between tissues becomes more evident when comparing the effects of the *clsy* double mutants, as significantly more reduced 24nt-siRNA clusters and hypo CHH DMRs are observed in the *clsy3,4* double mutant in rosette versus leaf tissues. For ovules, the genetic dependency is the opposite, as the *clsy3* and *clsy3,4* mutants, rather than the *clsy1* and *clsy1,2* mutants, have the largest numbers of reduced 24nt-siRNA clusters and CHH DMRs (**Fig. 3C** and **Fig. S8**). Notably, for ovules, and to a lesser extent flowers, the number of reduced 24nt-siRNA clusters and hypo CHH DMRs in the *clsy1* and *clsy4* mutants is higher than expected based on the 24nt-siRNA abundance data. Therefore, in addition to highly expressed 24nt-siRNA clusters controlled by CLSY3, more lowly expressed clusters controlled by CLSY1 and CLSY4 (**Fig. 3C** and **Fig. S8-9**) also play roles in shaping the 24nt-siRNA and DNA methylation landscape in both these tissues. Finally, across all these tissues, CLSY2 has a minimal effect (**Fig. 3B, C** and **Fig. S6-9**), suggesting that this CLSY may play a supporting role in general, or perhaps a more dominant role in tissues or cell types not yet profiled. Thus, depending on the tissue, different combinations of CLSY family members, or even individual CLSY proteins, control 24nt-siRNA and DNA methylation patterns at thousands of sites throughout the genome, establishing the CLSY proteins as tissue-specific regulators of the RdDM pathway.

Beyond the newly discovered tissue-specific roles for the CLSY family, comparisons between 24nt-siRNA and DNA methylation levels across the four tissues demonstrate that several behaviors initially observed in flowers^34^ appear to be conserved properties of the CLSY family. First, with the exception of leaf and rosette tissues that are dominated by the activity of CLSY1, clusters with reduced expression in the *clsy* single and double mutants are largely non-overlapping for all the other tissues (**Fig. 3C**). Furthermore, these reductions, especially for the double mutants, approach the levels observed in the *pol-iv* mutant and are less reduced, if at all, in the other *clsy* single mutants as compared to wild-type controls (**Fig. 3D** and **Fig. S6-9**). Together, these findings demonstrate that controlling 24nt-siRNAs in a locus-specific manner is a common feature of the CLSY proteins across tissues. Second, similar genetic relationships between CLSY family members were observed across all tissues: the *clsy* doubles displayed larger 24nt-siRNA reductions and affect more loci than their single mutant counterparts, and in all cases the *clsy* quadruple mutant essentially phenocopies *pol-iv* (**Fig. 3D** and **Fig. S6-9**), demonstrating that the CLSY family is responsible for controlling Pol-IV function across a diverse set of tissues. Third, across all tissues the *shh1* mutant phenocopies the *clsy1,2* double mutant, suggesting a conserved mechanism of targeting for these CLSYs irrespective of the tissue (**Fig. 3D, E**). Finally, for all combinations of *clsy* mutants, the loss of 24nt-siRNAs is associated with reduced CHH (and to a lesser extent CHG) methylation (**Fig. 3E** and **Fig. S6-9**). Collectively, these findings demonstrate that CLSYs act as locus-specific regulators of the RdDM pathway in diverse tissues and control tissue-specific patterns of DNA methylation in a manner consistent with their expression profiles during plant development.

### CLSY3 targets Pol-IV to siren loci to control 24nt-siRNA and DNA methylation levels

Given the dominant roles for CLSY3 and CLSY4 in ovules (**Fig. 3**), connections between these CLSYs and siren loci, which dominate the 24nt-siRNA landscape in ovules (**Fig. 2A**), were investigated. Based on 24nt-siRNA cluster overlaps, siren loci are almost exclusively controlled by CLSY3 and CLSY4. Of the 133 siren loci, 132 overlap with *clsy3,4*-dependent 24nt-siRNA clusters and 106 overlap with *clsy3*-dependent clusters in ovules (**Fig. 4A**). In contrast, none of the *clsy4*-dependent 24nt-siRNA clusters overlap with siren loci, demonstrating that although there is some redundancy with CLSY4, CLSY3 is the primary CLSY acting at these sites. Consistent with these overlaps, 24nt-siRNA levels at siren loci in ovules and flowers are specifically reduced in *clsy3, clsy4*, and *clsy3,4* mutants, with the *clsy3* single mutant showing stronger reductions than observed in the *clsy4* single mutant (**Fig. 4B**). These analyses also further illustrate the tissue-specific expression of these clusters as their levels are significantly lower in flower tissue and are barely detectable in leaf and rosette tissues (**Fig. 4B**). For methylation, the tissue-specificity is even more striking, as DNA methylation levels in all sequence contexts are reduced at siren loci in the *clsy3,4* double, *clsy* quadruple, and *pol-iv* mutants in ovules, but are already low in the other tissues and remain largely unaffected in RdDM mutants (**Fig. 4C** and **Fig. S10A**). Interestingly, only minimal reductions in DNA methylation are observed in the *clsy3* single mutant in ovules (**Fig. 4C**), suggesting that the 24nt-siRNAs remaining in this mutant are sufficient to maintain near wild-type levels of methylation. Although it cannot be excluded that some additional floral tissues produce 24nt-siRNAs at siren loci, these findings suggest that most, if not all, 24nt-siRNAs at these sites arise from ovules, that they are required to maintain DNA methylation in all sequence contexts, and that their production is most strongly dependent on CLSY3.

**Figure 4.**
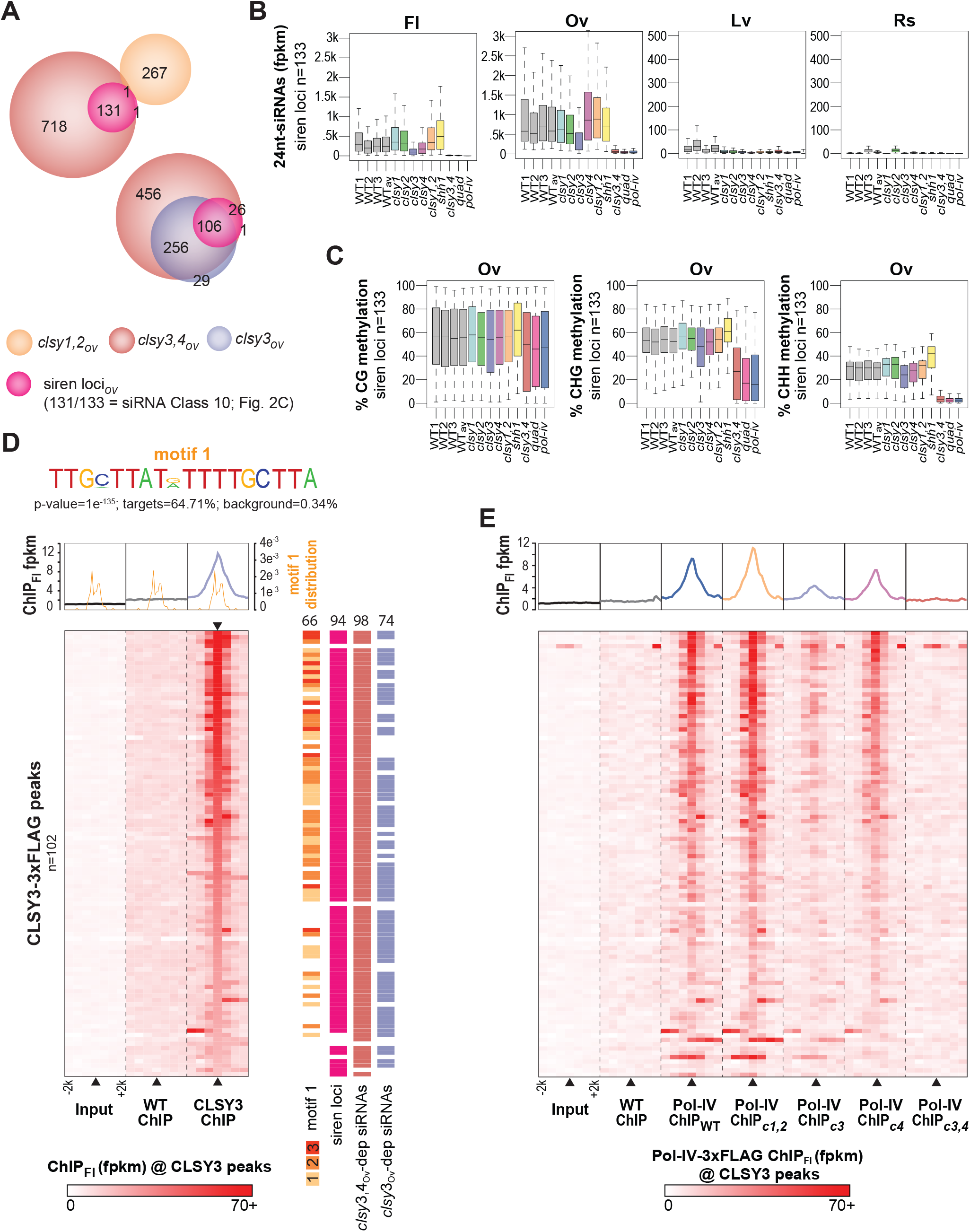
CLSY3 and CLSY4 are required for 24nt-siRNA production and Pol-IV recruitment at siren loci. (**A**) Scaled Venn diagrams showing the relationships between siren 24nt-siRNA clusters and those with reduced expression in the indicated *clsy* mutants from ovule tissue. siren loci are colored based on the 24nt-siRNA Class (Class 10) in which they reside based on **Fig. 2C**. (**B** and **C**) Boxplots showing 24nt-siRNA or DNA methylation levels at siren loci (n=133) in the indicated genotypes and tissues, respectively. (**D** and **E**) Metaplots (upper) and heatmaps (lower) showing the levels of CLSY3 enrichment (**D**) or Pol-IV enrichment (**E**) at CLSY3 ChIP-seq peaks (n=102) and surrounding genomic regions (+/-2kb). These heatmaps are all in the same order, based on a high to low ranking of CLSY3 enrichment in **C** as indicated by the inverted black triangle. For the Pol-IV ChIP data the IP subscripts indicate the genetic background (*e*.*g*. IP_c3_ indicates the *clsy3* background). In **D**, the distribution of motif 1 is included on a second y-axis scale and the occurrences of this motif, as well as the overlaps between the CLSY3 peaks and siren loci, *clsy3*-dependent, or *clsy3,4*-dependent 24nt-siRNA clusters are indicated to the right of the heatmap.

To determine whether CLSY3 acts directly at siren loci and whether it is required for Pol-IV recruitment to these sites, chromatin immunoprecipitation (ChIP)-seq experiments were conducted using flower tissue from wild-type (Col) plants or *clsy3* mutant plants expressing a tagged version of *CLSY3* under its endogenous promoter (CLSY3-3xFLAG, **Table S13**). 102 CLSY3 peaks were identified from the resulting ChIP-seq data (Col ChIP versus CLSY3-3xFLAG ChIP FC>2.5 and FDR< 0.001; **Table S14**), and were also enriched relative to an input control (**Fig. 4D**). Analyses of these CLSY3 peaks revealed the presence of a highly conserved motif centered under ∼2/3 of the peaks, with a higher proportion and number of motifs identified at sites with the most CLSY3 enrichment (**Fig. 4D** and **Fig. S10B**). In addition, these peaks were found to overlap with siren loci (94/102) as well as *clsy3-* and *clsy3,4*-dependent 24nt-siRNA clusters (74 and 98/102, respectively) (**Fig. 4D** and **Fig. S10C**). Finally, using previously published Pol-IV ChIP-seq data^34^, it was confirmed that CLSY3 and CLSY4, but not CLSY1 and CLSY2, are required for Pol-IV enrichment at these peaks (**Fig. 4E**). Together, these data demonstrate that CLSY3 is localized to siren loci, possibly through the direct or indirect recognition of a conserved sequence motif, and is required for the recruitment of Pol-IV to facilitate the production of very high levels of 24nt-siRNAs at siren loci specifically in ovules.

### Loci regulated by different CLSY pairs behave differently across and between tissues

To compare the behaviors of loci controlled by CLSY1 and CLSY2 versus CLSY3 and CLSY4, 24nt-siRNA clusters regulated by these pairs of CLSYs were identified across all tissues (*i*.*e. clsy1,2*- and *clsy3,4*-dependent_Fl,Rs,Lv,Ov_ 24nt-siRNA clusters, respectively), and separated into 8 groups each based on the average wild-type expression levels at these sites (**Fig. 5A, B**). Across the four tissues, a higher degree of variation was observed between groups for the *clsy3,4*-dependent_Fl,Rs,Lv,Ov_ clusters, which was further verified by calculating the coefficient of variation (CV) of these loci for all four tissues (Fl,Rs,Lv,Ov), as well as after excluding the ovule data (Fl,Rs,Lv; **Fig. 5C**). Assessment of CHH methylation levels at these same sets of 24nt-siRNA clusters revealed a similar pattern and again showed significantly more variation at sites controlled by CLSY3 and CLSY4 (**Fig. 5A-C**). In addition to variation in expression levels, the chromosomal distribution of *clsy3,4*-dependent_Fl,Rs,Lv,Ov_ clusters are also more diverse (**Fig. 5D, E** and **Fig. S11**). While *clsy1,2*-dependent_Fl,Rs,Lv,Ov_ 24nt-siRNA clusters are enriched in the chromosome arms across all tissues, the *clsy3,4*-dependent_Fl,Rs,Lv,Ov_ clusters shift from being enriched in pericentromeric heterochromatin in flower and rosette tissues to being more evenly distributed in ovules and leaves (**Fig. 5D, E** and **Fig. S11**). Finally, transcriptome profiling in the *clsy* mutants across tissues revealed that the *clsy4* single and *clsy3,4* double mutants reactivate the largest numbers of *pol-iv-*dependent genes and TEs (**Fig. 5F**). Although it remains to be determined why loci controlled by CLSY3 and CLSY4 vary more in their distribution and their effects on 24nt-siRNA levels, DNA methylation patterns, and TE silencing, these data demonstrate that the relatively smaller set of sites controlled by these factors play a disproportionately large role in shaping the epigenomes across the four tissues analyzed.

**Figure 5.**
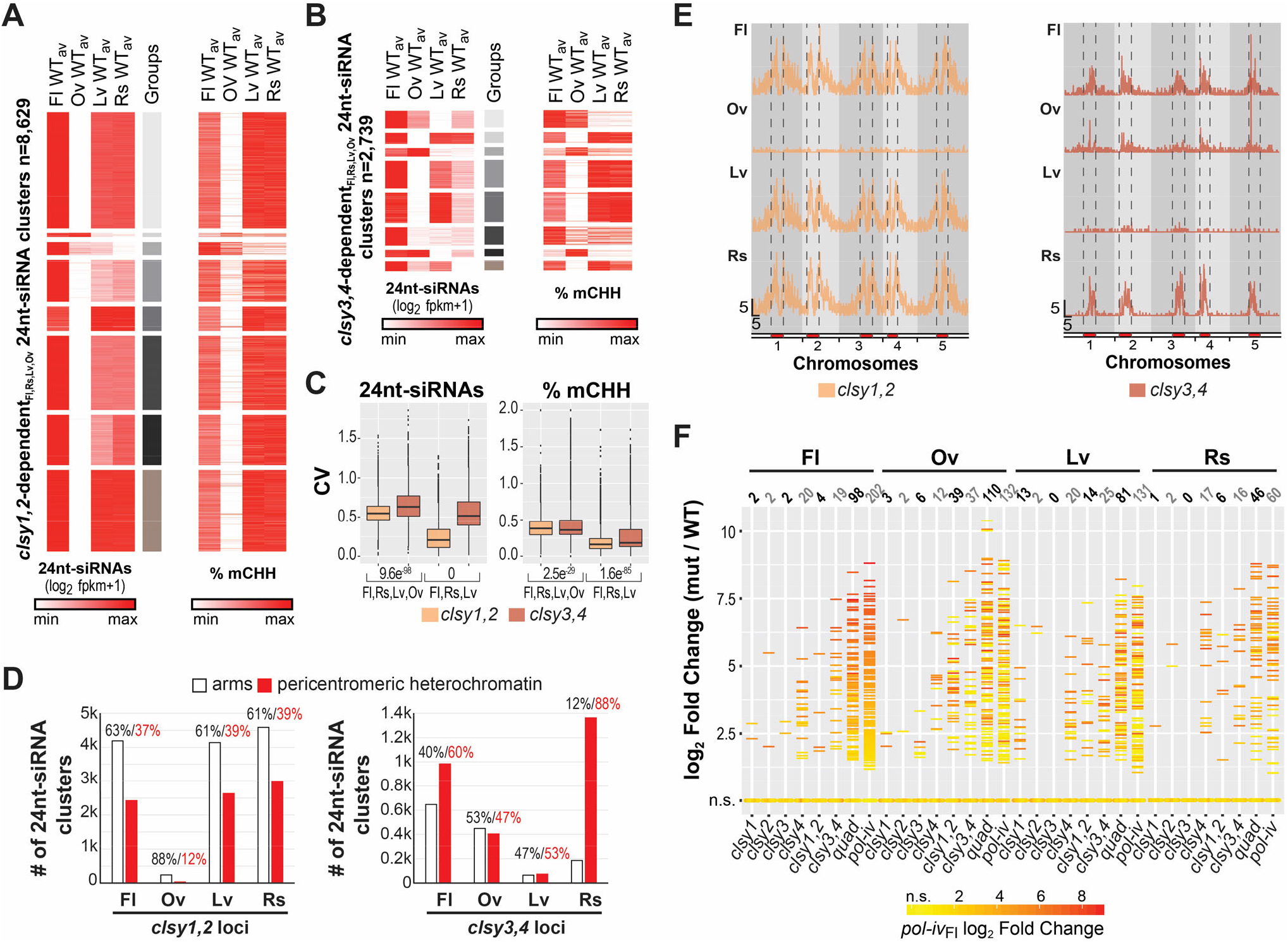
CLSY3 and CLSY4 play a disproportionately strong role in shaping the epigenome.

(**A** and **B**) Heatmaps showing the WT_av_ levels of 24nt-siRNAs and DNA methylation for each tissue at the combined set of 24nt-siRNA clusters reduced in either the *clsy1,2* (**A**) or *clsy3,4* (**B**) mutants. The heatmaps were clustered (grey scale) into 8 groups based on their 24nt-siRNA levels and the CHH methylation levels at these regions are shown in the same order to the right. (**C**) Boxplots showing the coefficient of variation (CV) between the 24nt-siRNA expression levels (log_2_ fpkm+1) or percent CHH methylation (% mCHH) at the regions shown in **A** and **B**. Comparisons were made including all four tissues (Fl, Rs, Lv, and Ov) or just three tissues (Ov excluded). P-values based on t-tests are indicated for each *clsy* mutant pair below each graph. (**D**) Bar graph showing the number of 24nt-siRNA clusters (1k=1000) dependent on *clsy1,2* or *clsy3,4* that reside in the chromosome arms versus pericentromeric heterochromatin, as designated in Yelina *et al*.^66^. The percent of clusters in each category is indicated above. (**E**) Distributions of reduced 24nt-siRNA clusters in the *clsy1,2* or *clsy3,4* mutants along the 5 chromosomes. The pericentromeric heterochromatin is marked in red and denoted by vertical dashed lines. Scale bars for x-axis (Mb) and y-axis (clusters/100kb bin) are indicated in the lower left corner and the tissue types are indicated in the upper left corner of each track. (**F**) Plot showing the expression levels [log_2_ FC (mut/WT)] of the 326 *pol-iv*-upregulated transcripts represented as horizontal lines (log_2_ FC ≥ 1.5 and FDR ≤ 0.05) or as dots (not significantly upregulated; n.s.) in the four tissues. The horizontal lines are colored based on the expression level in *pol-iv* from flower tissue and the number of DE loci are indicated above each column

To investigate the relative contributions of the CLSYs in shaping the differences in the 24nt-siRNA and DNA methylation landscapes between tissues, the effect of each *clsy* mutant combination on the profiles at 24nt-siRNA clusters and WTvsWT CHH DMRs were determined on a genome-wide scale for each tissue. For 24nt-siRNAs, PCA analysis (**Fig. 6A**) and assessment of expression levels at the master set of 24nt-siRNA clusters (**Fig. 6B**) revealed major shifts in RdDM mutants. For example, in flower tissue, *shh1* or *clsy1,2* mutants show profiles most similar to wild-type samples from ovules, while in ovule tissue, *clsy3,4* mutants show profiles most similar to wild-type samples from leaves (**Fig. 6A, B;** colored arrows). In leaf and rosette tissues, the *clsy1* singles already show similar profiles as *pol-iv* mutants, unlike in other tissues where only the *clsy* quadruple mutants behave like *pol-iv* mutants (**Fig. 6A, B;** grey arrows), further underscoring the dominant role for CLSY1 in these tissues. For DNA methylation, PCA analysis (**Fig. 6C**) and assessment of methylation levels in all three contexts (**Fig. 6D;** all mC) at the WTvsWT hypo CHH DMRs that overlap with the 10 24nt-siRNA classes (n=11,511/12,532) revealed similar shifts (**Fig. 6C, D**; grey and colored arrows). Here, the main difference between the behaviors of the 24nt-siRNAs and DNA methylation profiles occurred at 24nt-siRNA classes enriched for pericentromeric loci (classes 5-8) where fewer DMRs were identified (**Fig. 6B, D**), in line with the known roles of the CMT2/3 pathways in maintaining DNA methylation in these regions^27,28^. For both the 24nt-siRNA and DNA methylation analyses, the overall changes in the epigenetic profiles for each genotype and tissue are quantified by class in **Fig. S12**. Together, these findings demonstrate that disruption of specific *CLSY* family members is sufficient to shift the global 24nt-siRNA and, to a lesser extent DNA methylation, landscapes between tissue types (*e*.*g*. between flowers and ovules) or to largely phenocopy loss of Pol-IV (*e*.*g. clsy1* in leaves), revealing the extent to which these tissue-specific regulators contribute to the observed epigenetic diversity.

**Figure 6.**
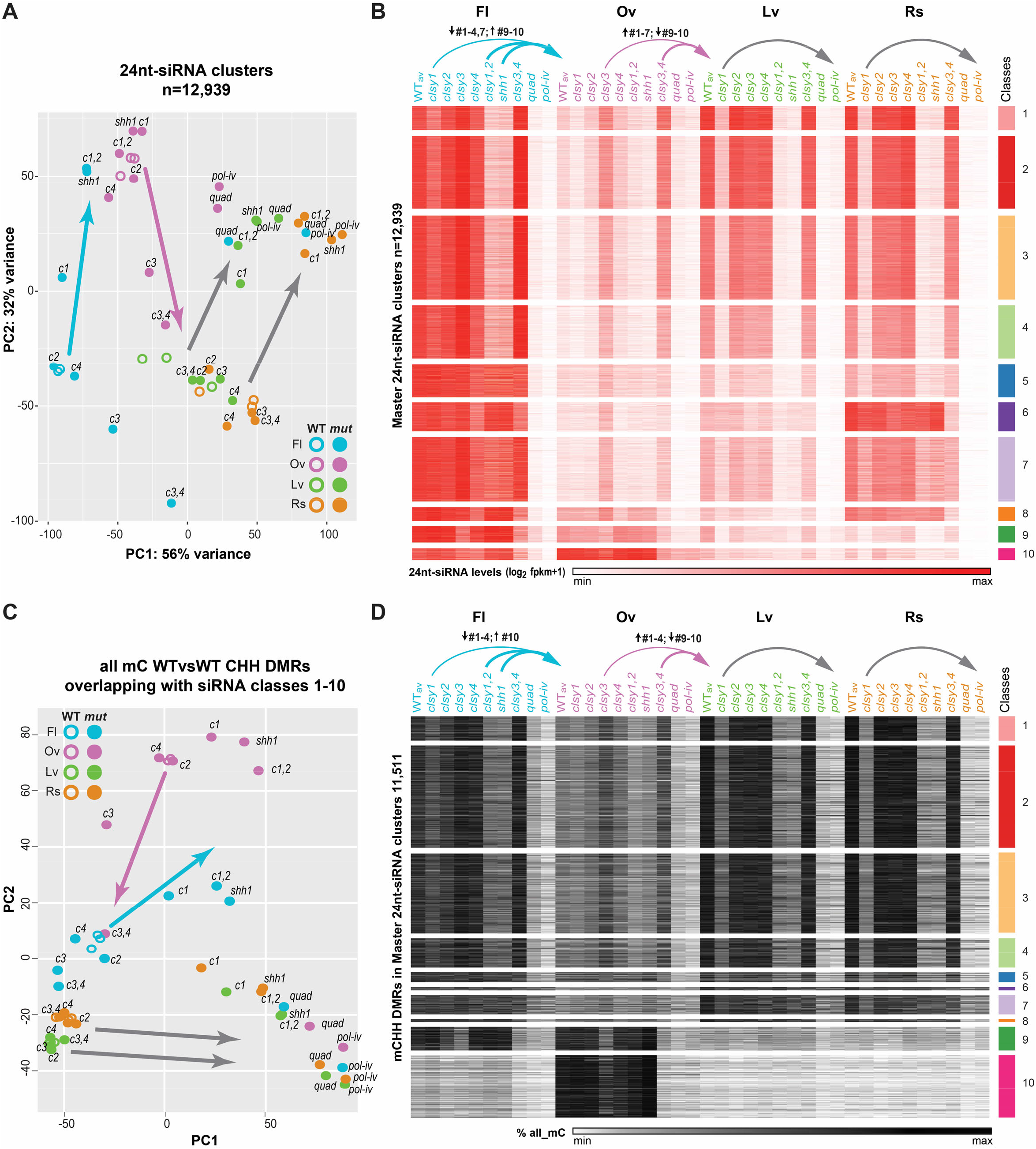
*clsy* mutants shift the epigenetic landscapes between tissues.

(**A** and **C**) Principal Component Analysis (PCA) plot of 24nt-siRNA or DNA methylation levels in all contexts (all mC) at the 12,939 master 24nt-siRNA clusters compiled from all four tissues (**Table S5**) or at the WTvsWT CHH DMRs (**Table S7**) that overlap with these clusters, respectively. WT samples (three replicates; open circles) and various RdDM mutants (filled circles) are colored by tissue, as indicated at the bottom right or top left. Mutants are abbreviated as follows, *c1=clsy1* etc. (**B** and **D**) Heatmaps showing 24nt-siRNA and DNA methylation levels in all sequence contexts (all mC) at reduced 24nt-siRNA clusters grouped by the ten classes defined in **Fig. 2C** (**B**) or at the WTvsWT CHH DMRs within these clusters (**D**) for the genotypes and tissues indicated above. The color scales and the numbers (n) of clusters or DMRs are as indicated on the x- and y-axes, respectively. In all panels, the arrows show mutants with patterns that resemble other WT tissues (colored) or single mutants that mimic *pol-iv* (grey). In **B** and **D**, the classes that are upregulated or downregulated are indicated above the colored arrows.

## Discussion

Tissue- and cell type-specific patterns of DNA methylation have been identified in increasing detail across diverse eukaryotic organisms. However, the cellular processes and machinery responsible for generating such patterns have remained largely elusive. Here, our findings demonstrate that through a combination of tissue-specific expression and locus-specific targeting, the CLSYs control 24nt-siRNAs and DNA methylation patterns during plant development. While the differences in the epigenomes profiled are not solely regulated by RdDM, this pathway is responsible for the vast majority of robust DNA methylation changes between tissues (**Fig. 2E** and **Fig. S3A**). Furthermore, our data links these changes to regulation of 24nt-siRNAs by the CLSY proteins (**Fig. 3** and **Fig. S6-9**), demonstrating they are major players shaping tissue-specific methylation patterns. Amongst the tissues profiled, we observed a range of different genetic dependencies on the CLSYs, with flower tissue showing the most even reliance across all four CLSYs, while ovules are most strongly dependent on CLSY3 and CLSY4, and leaf and rosette tissues are strongly dependent on CLSY1. These tissue-specific differences and large effects on the epigenome match the observed expression patterns of the CLSYs (**Fig. 1**) and are consistent with the observation that specific *clsy* mutants are sufficient to shift the epigenetic landscapes between tissues (**Fig. 6**), respectively. Taken together, these findings allow us to pair, for the first time, the distinct epigenetic profiles observed in flower, ovule, leaf and rosette tissues with the responsible cellular machinery (*i*.*e*. specific CLSY proteins), providing substantial mechanistic insights into the processes that enable the generation of diverse methylation patterns during plant development and addressing a largely unanswered question in the field of epigenetics.

Given the complexity of the profiled tissues, our findings likely under-represent both the cell type specificity and the contribution of the CLSYs in controlling DNA methylation patterns. For example, in the flower, leaf, and rosette tissues, our GUS staining experiments demonstrate that only a subset of cell types in each tissue express detectable levels of a given *CLSY* reporter (**Fig. 1B**). Based on these observations, we hypothesize that the CLSYs likely control 24nt-siRNA and DNA methylation patterns on an even finer scale than reported here. We further hypothesize that more cell type-specific profiling will also reveal larger DNA methylation decreases in the *clsy* mutants, since potentially unaffected methylation patterns in neighboring cells, which are included in our current data, would be excluded from the percent methylation ratios. Perhaps the exception to this notion of under-representation, is the data from ovules. Here we see more uniform GUS expression (**Fig. 1B**) and we see the best correlation between 24nt-siRNA and mCHH patterns on a global scale (**Fig. 2A** and **Fig. S2A**). In moving forward, it will be important to test the aforementioned hypotheses and to determine whether the high degree of correlation in ovules is specific to this tissue or whether such correlations will be observed in other instances where more uniform cell types are able to be profiled. Nonetheless, our current data already demonstrate that by combining the activities and expression levels of the CLSYs in different ways, a variety of 24nt-siRNA and DNA methylation patterns can be generated. Furthermore, recent studies profiling epigenetic changes during Arabidopsis embryogenesis^6^ and seed germination^8,38^ both show dynamic *CLSY* expression patterns, suggesting that modulation of just these four factors could be leveraged to generate a plethora of different patterns of DNA methylation throughout plant development.

Undoubtedly, one of the most unique patterns of 24nt-siRNAs and DNA methylation was observed in ovules. In this tissue, the small RNA landscape is dominated by a small number of loci that produced high levels of 24nt-siRNAs (**Fig. 2A, B** and **Fig. S2A**). This distinct pattern of 24nt-siRNAs was initially observed in the endosperm of rice, where they were designated as siren loci^36^. Subsequently, the small RNA patterns leading to this designation have been detected in other plant species and have been shown to arise prior to formation of the endosperm, in ovule tissues^37,39,40^. Regarding their function, our work in Arabidopsis (**Fig. 2A, Fig. S2A**, and **Fig. 4C**), and work from Grover *et al*. in *B. rapa*^37^, demonstrate that, at least in these species, siren 24nt-siRNAs target high levels of DNA methylation via the RdDM pathway. Thus, by this metric, they behave similarly to other 24nt-siRNAs. However, given their locations near genes^36,37^ and findings that mutations in Pol-IV components result in severe fertility defects in *B. rapa*^*41*^ and rice^42^, and subtle defects in Arabidopsis^41^, it has been suggested that siren loci could play roles in regulating genes important for seed development and/or for controlling imprinting in the endosperm^36,37^. Although examination of the 24nt-siRNA, DNA methylation, and gene expression profiles at siren loci using our Arabidopsis ovule data did not show any clear candidates in support of these hypotheses, such roles may be more evident in species with stronger fertility defects. Alternatively, these 24nt-siRNAs could play additional roles outside of targeting methylation, akin to other reproduction-specific small RNAs like piwi-interacting RNAs (piRNAs), some of which target DNA methylation at transposons to facilitate gene silencing, while others fail to map to transposons and play regulatory roles that remain poorly understood^43^.

Regardless of their mode of action, the connections between siren loci and seed development, a process essential for crop yields, underscore the importance of understanding how the production of 24nt-siRNAs at these loci are regulated. Here, our findings provide several new mechanistic insights: First, we demonstrate that 24nt-siRNA production at siren loci relies mainly on CLSY3, with some contributions by CLSY4, such that these 24nt-siRNAs are reduced to *pol-iv* levels in *clsy3,4* double mutants (**Fig. 4A, B**). Second, we demonstrate that CLSY3 and CLSY4 behave differently in ovule tissue compared to flower tissue, as siren loci are distributed quite evenly across the Arabidopsis chromosomes in ovules (**Fig. 2A** and **Fig. S2A**), while CLSY3 and CLSY4 targets in flower tissue are enriched in pericentromeric heterochromatin (**Fig. 5D, E** and **Fig. S11**). Finally, our CLSY3 ChIP-seq experiments led to the identification of a robust sequence motif at siren loci (**Fig. 4D**). While it remains to be tested whether this motif alone, or in combination with ovule-specific proteins or chromatin marks, is involved in recruitment of CLSY3 and Pol-IV to siren loci, or alternatively, if it plays a more downstream role, perhaps in the regulation of neighboring genes, these findings suggest both genetic and epigenetic information are important for properly regulating siren loci.

Taken together, our characterization of the CLSY proteins suggests that to understand how they control DNA methylation patterns, it will be important to not only understand how they target Pol-IV in a locus-specific manner, but also how the *CLSY* genes themselves are regulated during plant development. Regarding the former, our previous findings demonstrated that CLSY1 and CLSY2 target Pol-IV in connection with H3K9 methylation via SHH1, while CLSY3 and CLSY4 target Pol-IV in connection with CG methylation via unknown mechanisms^34^. Here, we find that in all four tissues profiled, *clsy1,2* mutants phenocopy *shh1* mutants (**Fig. 3D, E**), and control loci enriched in the chromosome arms (**Fig. 5D, E** and **Fig. S11**), suggesting a conserved mechanism of Pol-IV targeting by these CLSYs across diverse tissues. However, the different patterns of loci controlled by CLSY3 and CLSY4 in the four tissues (**Fig. 5**), combined with the presence of a motif at the siren loci that does not contain a CG dinucleotide (**Fig. 4D**), suggests these CLSYs may be targeted to chromatin via multiple pathways. If true, that would indicate that, within the CLSY family, we have identified both tissue-specific and tissue-independent mechanisms of epigenetic regulation. In contrast with these key insights into Pol-IV targeting, our understanding of how the *CLSY* genes are regulated during development, or possibly even in response to the environment, remains completely unknown and is a topic that will surely provide additional insights into the regulation of DNA methylation patterns in the future.

Finally, beyond striving to understand the roles of the CLSYs during normal plant development, our findings raise the possibility that the CLSYs could be leveraged to manipulate the epigenome in a tissue-specific manner and/or to better regulate DNA methylation patterns during tissue culture-based clonal propagation, a process that is known to alter the epigenome and generate epialleles that can negatively impact both plant health and crop yields^44,45^.

## Material and Methods

### Plant materials, growth conditions, and tissue collection

#### Plant materials

All plant materials used in this study were in the Columbia-0 (Col) ecotype. Previously characterized mutant lines include: *clsy1-7* (SALK_018319)^46,47^, *clsy2-2* (SAIL_484_F03)^34,48^, *clsy3-1* (SALK_040366)^34,47^, *clsy4-1* (SALK_003876)^34,47^, *nrpd1-4* (SALK_083051)^47,49^ and *shh1-1* (SALK_074540C)^32,47^.

#### Growth conditions and tissue collection

Growth conditions, tissue collection, and data for flower buds (Fl; stage 12 and younger) are described in Zhou *et al*.,^34^ (GEO accession: GSE99694). Growth conditions and tissue collection for 15 day old rosettes (Rs) and 25 day old leaves (Lv) for all genotypes were as follows: seeds were sterilized with bleach and plated on half strength Linsmaier and Skoog medium (1/2 LS; Caisson Labs, Cat# LSP03) supplemented with 1% agar. After three days stratification in the dark at 4 °C, plants were grown in a Percival growth chamber (GCU-36L5) under 16h-light and 8h-dark cycles at 22 °C for 7 days and were then transplanted onto soil in a Salk greenhouse with long day conditions. At day 15 and day 25 of total growth, rosette tissue (whole seedling without roots) or the first and second true leaves (without petioles) were collected from ∼45 plants as the rosette (Rs) and leaf (Lv) samples, respectively. These samples were immediately frozen in liquid nitrogen, and kept at -80 °C until use. Growth conditions and tissue collection for mature, unfertilized ovules (Ov) were as follows: seeds were grown on soil in a Salk greenhouse with long day conditions and un-opened flower buds (stage 10-12) were dissected using tweezers to obtain carpels. The carpels were then fixed on a glass slide with double-sided tape and opened using a straight sewing pin under a dissection microscope. The ovules were collected with the vacuum-based micro-aspiration device as detailed below. After collection, ovules for RNA extraction were quickly mixed with 100 µL Trizol (Invitrogen, Cat# 15596018), flash frozen in liquid nitrogen, and preserved at -80 °C until use, while ovules for DNA preparation were flash frozen in liquid nitrogen and preserved at -80 °C without Trizol. For each genotype, > 300 carpels were dissected and over 10,000 ovules were collected.

Ovules were collected in modified Eppendorf tubes using a vacuum-based micro-aspiration device engineered at the Salk Institute that consists of a collection tube and a vacuum pump as previously described in Sanchez-Leon, *et al*.^50^. The ovules were collected in a 1.5 mL microcentrifuge tube with the following lid modifications: (1) the hinge of the lid was cut and a 2 mm perforation as well as a 5 mm perforation were made in the center region of the lid with needles and scissors. (2) A ∼5 cm (15 µL capacity) glass capillary tube (Drummond Scientific, Cat# 1-1000-0150) was inserted in the 2 mm perforation such that the bottom of the tube reaches the 500 µl mark when the lid is refastened to the Eppendorf tube. (3) A Handy Plastic Tubing (external diameter 5.0 mm and inner diameter 3.5 mm (Momok, Cat# 20171004)) was inserted in the 5 mm perforation such that the bottom of the tubing reached the 1,000 µl mark and the bottom end was sealed with a nylon mesh filter using all purpose Krazy glue. (4) Perforations on the lid were sealed with glue sticks using a 100W Hot Glue Gun (Best, Cat# B073DCLKV4). (5) The modified lid was then reattached to the Eppendorf tube and placed inside a 15 mL conical falcon tube (without a cap) that is filled to the 12-13 mL mark with ground dry ice. (6) The top of the plastic tubing in the lid was attached tightly to a vacuum pump capable of ensuring a steady pressure of 0.3 to 0.6 Kg.cm^-2^ (the force necessary to detach an ovule from the placental tissue). After an ovule collecting session, the lid of the tube was carefully removed, and a clean new lid was used to cap the Eppendorf tube. To avoid contamination, lids with the capillary and plastic tubes were carefully washed to reuse or replaced with new lids for subsequent sessions.

#### Cloning, generation, and visualization of GUS reporters and tagged CLSY3 transgenic lines

For the GUS reporters, approximately 1.0-2.0 kb of the promoter regions from *CLSY1, CLSY2, CLSY3* and *CLSY4* were amplified from wild-type genomic DNA (**Table S15**) and cloned into the pDONR-P4P1R vector using the Gateway BP Clonase II Enzyme kit (Invitrogen, Cat# 11789020). These donor plasmids were recombined with a pDONR221 vector containing the *GUS* gene, a pDONR-P2RP3 vector with a mock fragment, and the pH7m34GW^51^ destination binary vector, using the Gateway LR Clonase II kit (Invitrogen, CA, Cat# 11791020). For the CLSY3 tagged line, a DNA fragment spanning the *CLSY3 gene*, including its endogenous promoter region, was amplified from wild-type genomic DNA (**Table S15**), and cloned into the pENTR/D-TOPO vector (Invitrogen, Cat# K240020). A carboxy-terminal 3x-FLAG-BLRP tag^32^ was inserted into a 3′ Asc I site present in the pENTR/D-TOPO vector. The pENTR/D construct was then digested with the MluI restriction enzyme (NEB, Cat# R0198S) and recombined into a modified gateway destination vector harboring Hygromycin drug resistance as described in Law *et al*. 2011^32^ using the Gateway LR Clonase kit (Invitrogen, CA, Cat# 11791019). The resulting constructs were sequence verified, transformed into the AGLO Agrobacterium tumefaciens strain, and used to create Arabidopsis transgenic lines in Col or *clsy3-1* mutant backgrounds for the GUS reporters or *pCLSY3+5’UTR::CLSY3-3xFLAG* lines, respectively, using floral dip method^52^. Primary transformants for each construct were selected on 1/2 LS medium with 25 mg/L Hygromycin and subsequent generations were screened to select for homozygous, single insert lines that were used for the GUS staining and ChIP experiments.

#### GUS histochemical staining of seedlings, leaves and flower buds

Rosettes, leaves, and flower buds from wild-type plants or plants harboring GUS reporters driven by the promoters of the four CLSYs (*pCLSY1-4::GUS*) were placed in ice-cold 90% acetone in water for 1 h, vacuum infiltrated for 10 min at room temperature, and then fixed for 20-30 min also at room temperature. These tissues were then briefly rinsed in GUS staining buffer (50 mM sodium phosphate buffer, pH 7.2, 0.2% Triton X-100, 2 mM Potassium ferrocyanide, and 2 mM Potassium ferricyanide) without X-Gluc and then vacuum infiltrated 2-3 times each for 10 min. The GUS staining buffer was then supplemented with fresh 2 mM X-Gluc (GoldBio, MO) and vacuum infiltrated again, 2-3 times each for 10 min. Samples were then incubated in the dark at 37 °C for ∼15 hours, except for the flower buds from the *pCLSY2::GUS* and *pCLSY4::GUS* lines which were incubated for 24 hours. The GUS enzymatic reaction was stopped and cleared by submerging tissues in a sequential series of ethanol washes (20, 35, 50 and 70% ethanol) for 30 min each. The cleared tissues were treated in chloral hydrate solution (1.5 mM in 30% glycerol), observed under a stereo microscope (Fisher, Cat# 03000015), and photographed with a SeBaCam14C digital camera (Fisher, Cat # SEBACAM14C CMOS 5V 14mp CMOS, Laxco) using SeBaView digital imaging software (Laxco).

### mRNA-seq library preparation, mapping, and data analysis

#### RNA isolation, mRNA-seq library construction, and sequencing

Total RNA from rosette and leaf tissue was isolated using the Quick-RNA MiniPrep kit (Zymo Research, Cat# R1055), while total RNA from ovules was isolated using the Direct-zol RNA Microprep Kits (Zymo Research, Cat# R2061). 1.0 μg total RNA (0.1-0.5 μg for ovules) was used to generate mRNA-seq libraries using the NEBNext Ultra II RNA Library Prep Kit (New England Biolabs, Cat# E7775). Sera-Mag Magnetic SpeedBeads (Thermo Scientific, Cat# 65152105050250) were used for all size selection and clean-up steps and the resulting libraries were pooled and sequenced (single end 50 bp, SE50) on a HiSeq 2500 machine (Illumina).

#### mRNA-seq mapping

mRNA-seq reads were mapped to the TAIR10 reference genome using STAR (v2.5.0c)^53^ with the following options: “--outFilterMismatchNmax 2”, to allow 2 mismatches, and “--outFilterMultimapNmax 1”, to include only uniquely mapped reads (**Table S1**). The sorted bam files were then used to generate Tag Directories using the (Hypergeometric Optimization of Motif EnRichment) HOMER^54^ makeTagDirectory script with the “-mis 2 and -unique” options and UCSC format tracks were generated using the makeUCSCfile script with the “tair10 -fragLength given and -style rnaseq” options.

#### Quantification of expression levels

To identify differentially expressed loci in each tissue and compare the expression of specific genes across tissues, raw read counts or normalized values for each sample were obtained using the HOMER^54^ analyzeRepeats.pl script with the “-noadj” or “-fpkm” options, respectively, and a custom .gtf file (**Source Data 1**). The .gtf file was generated based on the Araport11 annotation^55^ and includes genes, transposons/repeats, as well as several novel transcripts that were characterized in Zhou *et al*.^34^. The resulting expression files were then filtered to report just the isoform with highest expression across all genotypes (condense_genes_by_highest_sum_exp_header.py; **Source Data 1**). The expression patterns [log_2_ (fpkm +1)] of genes required for DNA methylation were visualized for representative wild-type samples from each tissue using MORPHEUS (https://software.broadinstitute.org/morpheus) (**Fig. 1C**). Differentially expressed loci for each tissue (log_2_ fold change (FC) ≥ 1 and FDR ≤ 0.05) were determined based on the raw read counts in R using Rstudio with the DESeq2^56^ package and default parameters including batch corrections for the ovule and rosette datasets that included samples from two different experiments.

#### Analysis of *pol-iv*-dependent upregulated loci across tissues

From the *pol-iv* upregulated loci in the four tissues, a list of 326 non-overlapping *pol-iv* upregulated loci was generated (*e*.*g*. if a gene and a TE annotation overlap, the most highly upregulated locus was selected) (**Table S10**). Of these 326 loci, those with significant upregulation (log_2_ FC ≥ 1 and FDR ≤ 0.05) in *pol-iv* mutants from each tissue (**Fig. 2G**) or (log_2_ FC ≥ 1.5 and FDR ≤ 0.05) across all the remaining mutants (**Fig. 5F** and **Table S9**) were visualized in R using RStudio with the following parameters: the matrix was organized by tidyr (gather), color-coded using tidyr based on the FC value in the *pol-iv* data from flowers, and then visualized by ggplot2 (geom_point). Loci outside the FC and FDR thresholds for each tissue were included as zero values in the matrix. The *pol-iv* dependent changes in epigenetic features surrounding these loci were plotted using deepTools (v2.4.0)^57^ (**Fig. S4A**) as described in Zhou *et al*.^34^. Briefly, mRNA-seq data sets were compared to wild-type controls with the bamCompare tool using the “--ratio=log2 --scaleFactorsMethod SES and -bs 10” options. smRNA-seq and MethylC-seq data sets were converted from wiggle (.wig) to bigwig format using the bedGraphToBigWig script and then compared to wild-type controls with the bigwigCompare tool using the “--ratio=log2 and -bs 10” options or with the bigwigCompare tool using the “--ratio=subtract” options, respectively. The resulting bigwig files were used to calculate a matrix with the computeMatrix tool using the “scale-regions -a 2000 --regionBodyLength 2000 -b 2000 and -bs=100” options and the data was plotted using the plotHeatmap tool and sorted based on the 24nt-siRNA values.

### ChIP-seq library preparation, mapping, and data analysis

#### Chromatin immunoprecipitation, library construction, and sequencing

A Flag-tagged CLSY3 line, pCLSY3::CLSY3-3xFlag in the *clsy3-1* mutant background, and non-transgenic wild-type plants were used for ChIP experiments. The ChIP was performed as previously described in Zhou *et al*.^34^. Briefly, for each genotype, ∼2.0 g of un-opened flower buds (stage 12 and younger) was collected, ground in liquid nitrogen, and in vitro crosslinked with 1% formaldehyde (20 minutes at RT; Sigma, Cat# F8775). The chromatin was then fragmented to ∼500bp by sonication, incubated with anti-Flag M2 Magnetic beads (4 °C for 2 h; Sigma, Cat# M8823), washed 5 times and then eluted with 3xFLAG peptide ([0.1 mg/mL]; Sigma, Cat# F4799). The crosslinking was reversed overnight at 65 °C and purified DNA (Thermo Scientific, Cat# 17908) was used to generate libraries with the NEBNext Ultra II DNA Library Prep Kit (New England Biolabs, Cat# 7645). The resulting libraries were sequenced (single end 50 bp, SE50) on a HiSeq 2500 machine (Illumina).

#### ChIP-seq data mapping, peak calling, and data analysis

The ChIP-seq data was aligned to TAIR10 reference genome using bowtie (v1.1.0)^58^ and the “-m 1 -v 2 --all –best and –strata” options to allow 2 mismatches and including only uniquely mapping reads (**Table S13**). TagDirectories and UCSC genome browser tracks from ChIP-seq data were generated with the makeTagDirectory and makeUCSCfile scripts from HOMER^54^ using the “format sam -mis 2 and -unique” or “none -fragLength given and -norm 10000000” options, respectively. The 102 CLSY3 ChIP peaks were identified with the HOMER^54^ findPeaks script using the “-style factor -region -L 2.5 -F 2.5 and -center” options (**Table S14**). Overlaps between these peaks and siren loci, *clsy3*-, and *clsy3,4*-dependent 24nt-siRNA clusters were determined using BEDOPS^59^ with the -e 1 option (**Fig. 4A**). Motifs within CLSY3 peaks (**Fig. 4D** and **Fig. S10B**) were identified using the findMotifsGenome.pl script from HOMER^54^ with the “-size given -len 6,8,10,12,18,20 -p 10 -mis 4 -bits” options and a bed file showing the positions of these motifs and the number of motifs per peak was generated using the annotatePeaks.pl script from HOMER^54^ with the “tair10 -m -mbed -matrix -nmotifs -multi” options (**Table S14**). Metaplots showing enrichment of reads from ChIP-seq datasets and the distribution of motif 1 over the CLSY3 peaks (**Fig. 4D, E**) were generated using the annotatePeaks.pl script from HOMER^54^ with the “tair10 -m -mbed -size 4000 -hist 100 -fpkm” options and plotted in Excel. Heatmaps showing enrichment of reads from ChIP-seq datasets (**Fig. 4D, E**) were generated using the annotatePeaks.pl script from HOMER^54^ with the “none -size 4000 -hist 600 -ghist -fpkm” options and plotted in Morpheus. The Pol-IV ChIP-seq data sets in different *clsy* mutants were downloaded from GSE99693^34^, and were mapped, analyzed, and visualized as described for the CLSY ChIP.

### smRNA-seq library preparation, mapping, and cluster calling

#### RNA isolation, smRNA-seq library construction, and sequencing

Total RNA extraction and small RNA enrichment were performed as previously described in Zhou *et al*.^34^. Briefly, 2.0 µg of total RNA from rosette and leaf tissues, or ∼0.5 µg of total RNA from ovules, were used for smRNA enrichment. The resulting small RNAs were used for library preparation with the NEBnext Multiplex Small RNA Library Prep Set for Illumina (New England Biolabs, Cat# E7300) following the user’s manual. The final library products were further purified using an 8% polyacrylamide gel to excise 130-160nt products relative to the pBR322 DNA-MspI Digest ladder (New England Biolabs, Cat# E7323AA). The purified libraries were pooled and sequenced (single end 50 bp, SE50) on a HiSeq 2500 machine (Illumina).

#### smRNA-seq mapping and 24nt-siRNA cluster calling

The smRNA-seq data were mapped (**Table S2**) and 24nt-siRNA clusters identified (**Table S4**) as described in Zhou *et al*.^34^. Briefly, the adapters were removed from the de-multiplexed reads using cutadapt (v1.9.1)^60^ and trimmed reads longer that 15nt were mapped to the TAIR10 genome using ShortStack (v3.8.1)^61^ with the following options: --mismatches 1 and either --mmap f, to include multi-mapping reads, or --mmap n, to include only unique mapping reads. As described in Zhou *et al*.^34^ only perfectly matched reads or reads with a single 3’ mismatch were used to call smRNA clusters using ShortStack^61^ with the --mincov 20, --pad 100, --dicermin 21 and --dicermax 24 options, to generate Tag Directories using the using the makeTagDirectory script from the HOMER^54^ with the -format sam -mis 1 and -keepAll options, and to make genome browser tracks specifically for the 24nt sized smRNAs using the makeUCSCfile script from HOMER^54^ with the -fragLength 24 and -norm 10000000 options.

#### Core, Master, and *clsy1,2*-versus *clsy3,4*-dependent_Fl,Rs,Lv,Ov_ 24nt-siRNA cluster lists (Table S5)

An inclusive list of 24nt-siRNA clusters present amongst the four tissues was generated by first identifying the core set of 24nt-siRNA clusters present in all three replicates for each tissue using the mergePeaks script from HOMER^54^, as was done for flower tissue in Zhou *et al*.^34^. These regions were then combined using the BEDOPS^59^ --merge function to identify a non-redundant set of clusters based on all four tissues, hereafter referred to as the “master 24nt-siRNA clusters” list. Lists of clusters reduced in the *clsy1,2* or *clsy3,4* mutants across all tissues were identified first using the BEDOPS^59^ -u function with the clusters from each of the four tissues as input files, and then the sort -u function to remove duplicates.

#### Differentially expressed (DE) 24nt-siRNA clusters analysis

For differential expression analyses, 24nt-siRNA levels at each of the master 24nt-siRNA clusters were quantified for each tissue and genotype with the HOMER^54^ annotatePeaks.pl script using the “-noadj, -size given and -len 1” options and differentially expressed 24nt-siRNA clusters were identified using DESeq2^56^ (FC ≥ 2 and FDR ≤0.01) (**Table S11**). To account for the extreme decreases in 24nt-siRNAs counts in some mutants (*i*.*e. pol-iv*), the size factors estimates calculated by DESeq2^56^ for the wild-type samples from each tissue were used to adjust the size factor estimates for the other samples to normalize the data to mapped reads of all size classes rather than to just 24nt-siRNAs. For details, see Zhou *et al*.^34^.

For each mutant in each tissue, a list of 24nt-siRNA clusters that are either differentially expressed or overlapped with one or more *pol-iv*-dependent hypo CHH DMRs (see “**DMR calling**”) was generated as follows: first the BEDOPS^59^ -n 1 function was used to find hypo CHH DMRs that do not overlap with DE clusters in the given mutant for each tissue and then the BEDOPS^59^ -e 1 function was used to identify clusters from the master 24nt-siRNA clusters list overlapping the DMRs. These clusters were then filtered to only include those showing a log_2_ FC < -0.415 (∼25% decrease), and finally the resulting 24nt-siRNA cluster list was combined with the DE clusters identified from the DEseq2^56^ analyses (**Table S11**). 24nt-siRNA levels at the clusters included based on their overlaps with hypo CHH DMRs were determined with the HOMER^54^ annotatePeaks.pl script using the “tair10 -size given -fpkm and -len 1” options (**Fig. S5A**).

### MethylC-seq library preparation, mapping, and Differentially Methylated Region (DMR) calling

#### DNA isolation, methyl-seq library construction, and sequencing

**∼**0.1 g of tissue from 15 day old rosettes, 25 day old leaves, or >5,000 unfertilized ovules were collected for genomic DNA isolation using the Dneasy Plant Mini Kit (Qiagen, Cat# 69104). MethylC-seq libraries were generated using 2.0 μg of DNA from rosettes and leaves, and 0.5-1.0 μg of DNA from ovules as described in Li *et al*.^62^. The libraries were then pooled and sequenced (single end 50 bp, SE50) on a HiSeq 2500 machine (Illumina).

#### MethylC-seq data processing

MethylC-seq reads were mapped and processed using BS-Seeker2^63^ as described in Zhou *et al*.^34^. Briefly, reads were mapped with the bs_seeker2-align.py script allowing 2 mismatches, clonal reads were removed using the MarkDuplicates function within picard tools (http://broadinstitute.github.io/picard/), and methylation levels at each cytosine covered by at least 4 reads were determined using the bs_seeker2-call_methylation.py script. For visualization of methylation levels in each sequence context or in all contexts together, wig files were generated using a custom perl script (BSseeker2_2_wiggleV2_CG_CHG_CHH_allmC.pl) based on the BS-Seeker2 CGmap output files. Information regarding the mapability, coverage, global percent CG, CHG, and CHH methylation levels, and non-conversion rates are presented in **Table S3**.

#### DMR calling

DMRs were determined as described in Zhou *et al*.,^34^. Briefly, DMRs in the CG, CHG or CHH contexts were identified in 100 bp non-overlapping bins based on either pair-wise comparisons between each mutant and three independent wild-type data sets, or between all three wild-type data sets from one tissue and all three wild-type datasets from another tissue. To be designated a DMR, the following criteria were required: (1) the 100 bp bins must to contain ≥ 4 cytosines in the specified context, (2) the bins must have sufficient coverage in both genotypes being compared (*i*.*e*. ≥ 4 reads over the required 4 cytosines in the specific context), and (3) the bins must show a fold change of 40%, 20% or 10% methylation in the CG, CHG, and CHH contexts, respectively, with an adjusted *p* value of ≤0.01. After these pairwise DMR calling, the final set of DMRs were determined for each mutant by taking the overlap between the DMRs called relative to each wild-type control (**Table S12**). For the DMRs between tissues, a more stringent approach was taken, requiring the DMR to be present in across all 9 pair-wise comparisons to be included (*e*.*g*. “WT_Rs_CG_hyperDMR_vs_Fl” **Table S7**). These wild-type (WT) versus WT DMRs were subsequently combined to generate a set of 100 bp hyper and hypo DMRs amongst all pairwise tissue comparisons (*e*.*g*. “WTvsWT all tissues hypo” DMRs; **Table S7** and **Fig. S3A**).

### 24nt-siRNA and DNA methylation data intersections, data analyses and visualization

#### Chromosome scale analysis of 24nt-siRNA and DNA methylation patterns

The levels of 24nt-siRNAs or CHH methylation for the three wild-type controls and *pol-iv* mutants were determined for each tissue in 5kb bins across the five Arabidopsis chromosomes with the HOMER^54^ annotatePeaks.pl script using ether the “tair10 -size given -len 1 and -ratio” or the “tair10 -size given -len 1 and -fpkm” options, respectively. For 24nt-siRNAs, a pseudo count of one was added to all values prior to a log_2_ transformation. After averaging the wild-type data, the 24nt-siRNA and methylation values were visualized as barplots in R using RStudio with the circularize package^64^ (**Fig. 2A** and **Fig. S2A**). For CHH methylation, between 1 and 3 bins, depending on the tissue, had values over 0.3%, but were capped at this value to facilitate visualization on a chromosome-wide scale.

#### PCA analyses

For **Fig. 6A** (PCA), raw values for 24nt-siRNA levels at the master clusters in the wild-type controls and *pol-iv* mutants for each tissue were calculated with the HOMER^54^ annotatePeaks.pl script using the “-noadj, -size given and -len 1” options and the PCA was plotted in R using RStudio with the DEseq2^56^ and ggplot2 packages. For **Fig. 6C**, the WTvsWT all tissues hypo CHH DMRs (**Table S7**) were filtered to include only DMRs overlapping with the 10 24nt-siRNA classes (**Table S6**), resulting in 11,511/12,532 DMRs. For all cytosines (CG, CHG, and CHH) within these DMRs, the coverage and frequencies of C and T bases were calculated from the ATCGmap output files from BS-seeker2 for each genotype. This information was then formatted for use with methylkit and plotted in R using RStudio with the methylkit package^65^ default options and a sd. threshold of 0.9.

#### Cumulative sum analysis

The cumulative sums of 24nt-siRNA levels were determined from the average values of the three wild-type replicates for each tissue at the master set of 24nt-siRNA clusters with the HOMER^54^ annotatePeaks.pl script using the “tair10 -size given -len 1 and -fpkm” options (**Fig. 2B**). The top 133 clusters, which correspond to 80% of all 24nt-siRNAs, were designated as siren loci and are presented in **Table S5**.

#### Chromosomal distributions

To visualize the chromosomal distributions of 24nt-siRNA clusters the TAIR10 genome was split into 100 kb bins and the number of 24nt-siRNA clusters from each class (1-10; **Fig. 2D**) or genotype (**Fig. 5E** and **Fig. S11A**) in each bin was determined using the bedmap^59^ count function requiring 1 bp overlap (--bp-ovr 1) and plotted along the five chromosomes with the pericentromeric regions as designated in Yelina *et al*.^66^. For the *clsy* mutants, the number and percent of clusters in each genomic region was also plotted in excel (**Fig. 5D**).

#### Pie Charts

For **Fig. 2E** the fraction of DMRs (**Table S12**) overlapping 24nt-siRNA clusters for each of the 10 classes (**Table S6**), or none, were determined with BEDOPS^59^ using the “-e 1” option. For **Fig. 3B**, the fraction of total 24nt-siRNAs present in specific categories were determined from raw read counts of a wild-type control from each tissue with the HOMER^54^ annotatePeaks.pl script using the “tair10 -size given -noadj and -len 1” options.

#### Venn Diagrams

Scaled Venn diagrams showing the overlaps between reduced 24nt-siRNA clusters (**Table S11**) or hypo CHH DMRs (**Table S12**) for the different sets of *clsy* single, double, and quadruple mutants were generated using VennMaster (version 0.38.2)^67^ (**Fig. 3C** and **Fig. S6-9**). Scaled Venn diagrams showing the overlaps between siren loci (**Table S5**), reduced 24nt-siRNA clusters (**Table S11**), and CLSY3 chIP peaks (BEDOPS^59^ “-e 1” option; **Table S14**) were also generated using VennMaster^67^. Unscaled Venn diagrams for 24nt-siRNA clusters (**Fig. S5B**) were generated with the same data using venny (https://bioinfogp.cnb.csic.es/tools/venny/index.html).

#### Coefficient of variation

To assess the degree of variation in the levels of 24nt-siRNAs and CHH methylation between *clsy1,2*- and *clsy3,4*-dependent_Fl,Rs,Lv,Ov_ 24nt-siRNA clusters across tissues, the coefficient of variation (cv= standard deviation average for each cluster across the 4 tissues) was calculated and plotted as boxplots in R using RStudio and p-values were calculated using t-tests (**Fig. 5C**).

#### Boxplots

For all boxplots showing the expression levels of 24nt-siRNAs (**Fig. 4B, Fig. S5A, Fig. S6-9B**, and **Fig. S12A**), the data was generated with the HOMER^54^ annotatePeaks.pl script with the “tair10 -size given -fpkm and -len 1” options and plotted in R using RStudio. For **Fig. S12A**, the resulting data was then log_2_ transformed after adding a pseudo count of one to each value. For all boxplots showing the percent methylation levels (**Fig. 4C, Fig. S6-9B, D, Fig. S10A** and **Fig. S12A**), the data was generated with the HOMER^54^ annotatePeaks.pl script using the “tair10 -size given -len 1 and -ratio” options and plotted in in R using RStudio.

#### Heatmaps and data clustering

For all heatmaps showing the expression levels of 24nt-siRNAs (**Fig. 2C, Fig. 3D, Fig. 5A, B**, and **Fig. 6B**) the data was generated with the HOMER^54^ annotatePeaks.pl script using the “tair10 -size given -fpkm and -len 1” options and plotted in Morpheus using the “log_2_+1” adjustment with the coloring set as “relative color scheme” based on the min and max values for each row. For **Fig. 2C** the data was split into 10 classes (**Table S6**) using the Morpheus “KMeans Clustering” tool by selecting the “cluster by rows” and “one minus Pearson correlation” options. For **Fig. 3D**, the data was ordered based on the overlaps in the reduced 24nt-siRNAs clusters (first for the singles, then doubles, then quadruple mutant clusters) and ranked high to low for each grouping (*e*.*g*. Fl *clsy1*-dependent clusters are ranked high to low based on the expression in the *clsy1* mutant). For **Fig. 5A** and **B**, the data was split into 8 groups using the Morpheus “KMeans Clustering” tool by selecting the “cluster by rows” and “one minus Pearson correlation” options.

For all heatmaps showing percent methylation levels (**Fig. 3E, Fig. 6D**, and **Fig. S3A**), the data was generated with the HOMER^54^ annotatePeaks.pl script using the “tair10 -size given -len 1 and -ratio options” and plotted in Morpheus with the coloring set as “relative color scheme” based on the min and max values for each row for **Fig. 3E** and **Fig. 6D**, or as a max percent methylation for **Fig. S3A**. In **Fig. 3E** the heatmap is ordered exactly as in **Fig. 3D** and for **Fig. 6D** the DMRs are ordered based on the 24nt-siRNA classes they overlap with. In **Fig. S3A**, the data was split into 8, 7, and 2 classes for the CHH, CHG, and CG contexts, respectively, using the Morpheus “KMeans Clustering” tool by selecting the “cluster by rows” and “one minus Pearson correlation” options (**Table S8**).

### Data availability

Illumina sequencing data (smRNA-seq, MethylC-seq, mRNA-seq, and ChIP-seq) has been deposited in the NCBI Gene Expression Omnibus (GEO) and are accessible through the GEO series accession number GSE165001.

## Supporting information

Table S1. Summary of mRNA-seq data

Table S2. Summary of smRNA-seq data

Table S3. Summary of MethylC-seq data

Table S4. 21-24nt small RNA clusters

Table S5. 24nt-siRNA cluster lists and summary

Table S6. 24nt-siRNA heatmap clustering and summary

Table S7. DMRs between WT tissues and summary

Table S8. DNA methylation heatmap clustering and summary

Table S9. clsy upregulated DEGs overlapping with pol-iv and summary

Table S10. pol-iv DE analysis and summary

Table S11. DE 24nt-siRNA clusters and summary

Table S12. hypo DMRs and summary

Table S13. Summary of ChIP-seq data

Table S14. ChIP analysis and summary

Table S15. Primers table

GTF file

## Figure Legends

**Figure S1. Visualization of *CLSY* expression patterns using ePlant**^35^. Relative expression levels of the four *CLSY* genes from two Plant eFP viewers: (**A**) AtGenExpress^69,70^ and (**B**) RNA-seq^68^. For comparison, the color scales are equal for panels **A** and **B** with the exception of *CLSY3*, which is on a higher scale, as indicated by the red text on the color bar. All tissues and organs are labeled as in the ePlant viewers^35^. Images are not to scale.

**Figure S2. Global characterization of 24nt-siRNA and CHH methylation patterns across tissues**. (**A**) Circular genome view showing the patterns of 24nt-siRNAs (light grey background) and CHH methylation (dark grey background) across all five chromosomes (chr1-5) in 5kb bins based on the WT_av_ expression levels from each tissue. The color scales for the data sets are as indicated in the center of the plot and the tracks are labeled every 5Mbs, with the pericentromeric heterochromatin, as designated in Yelina *et al*.^66^, marked in red along the outer circle. For CHH methylation, between 1 and 3 bins, depending on the tissue, had values over 0.3 (0.3^+^), but were capped at this value to facilitate visualization on a genomic scale.

**Figure S3. Tissue specific comparison of DNA methylation patterns**. (**A**) Scaled heatmaps showing the designation of 8 CHH, 7 CHG, and 2 GC classes of hypo DMRs, respectively, based on the methylation levels (WT_av_ and *pol-iv*) for each tissue at the master set of WTvsWT DMRs for each context (**Table S7**). Classes showing clear reductions in methylation in the *pol-iv* mutant are indicated as *pol-iv*-dependent. For CG DMRs, this is indicated by an asterisk.

**Figure S4. Assessment of 24nt-siRNA and DNA methylation levels surrounding differentially expressed transcripts**. (**A**) Heatmaps and profile plots showing the expression of all *pol-iv* upregulated transcripts (log_2_ FC ≥ 1 and FDR ≤ 0.05; as shown in (**Fig. 2G**)) as well as the corresponding 24nt-siRNA and DNA methylation levels at these same loci. For the mRNA and 24nt-siRNA analyses, the log_2_ fold changes in expression in *pol-iv* mutants relative to wild-type controls are plotted and for the DNA methylation analysis, the difference in the percent methylation between *pol-iv* mutants and wild-type controls is plotted. Color bars indicating the scales are shown below. The heatmaps include 2kb flanking the transcription start site (S) and the transcription termination site (T) and were ranked based on the 24nt-siRNA and mCHH values for each tissue. The profiles for the genes or TE/repeats are shown in blue and light green, respectively, above each heatmap. For flower tissue, the data from Zhou *et al*.^34^ was reanalyzed using an updated genome annotation (**Source Data 1**).

**Figure S5. Additional analysis of 24nt-siRNA clusters affected in the different tissues and mutant backgrounds**. (**A**) Boxplots showing 24nt-siRNA levels at clusters for each tissue and genotype that overlap with hypo CHH DMR(s) and are reduced (>25%) compared to tissue matched WT controls, but were not initially detected as DE clusters due to the FC and p-value thresholds. Here, and for all subsequent boxplots, the graphs show the interquartile range (IQR), with the median shown as a black line and the whiskers corresponding to 1.5 times the IQR. Above each plot, the numbers (n) of clusters are indicated and biological replicates for the WT controls are designated as WT1, WT2, and WT3, with the average signal from these replicates designated as the WT_av_. (**B**) Unscaled Venn diagrams showing the overlaps between 24nt-siRNA clusters affected in each mutant combination and tissue type. Mutants are colored as indicated on the right. * indicates overlaps not included in the scaled Venn diagrams due to spatial constraints.

**Figure S6. Relationships between 24nt-siRNA and DNA methylation levels for *clsy* mutants in rosette (Rs) tissue**. (**A** and **C**) Scaled Venn diagrams based on the reduced 24nt-siRNA clusters provided in **Table S11** or the hypo CHH DMRs provided in **Table S12** showing the relationships between loci affected in the indicated single, double, and quadruple mutants relative to *pol-iv*. For readability, only overlaps >20 are labeled. A small number of overlaps are not shown due to spatial constraints, but unscaled Venn diagrams showing all the overlaps are present in **Fig. S5B**. For the 24nt-siRNA clusters, the Venn diagrams for the single and double mutants are also shown in **Fig. 3C**. (**B** and **D**) Boxplots showing 24nt-siRNA and DNA methylation levels at reduced 24nt-siRNA clusters (**B**) or showing DNA methylation levels at hypo CHH and CHG DMRs (**D**). Above each plot, the numbers (n) of reduced 24nt-siRNA clusters identified for each mutant are indicated and biological replicates for the WT controls are designated as WT1, WT2, and WT3, with the average signal from these replicates designated as the WT_av_.

**Figure S7. Relationships between 24nt-siRNA and DNA methylation levels for *clsy* mutants in leaf (Lv) tissue**. (**A** and **C**) Scaled Venn diagrams based on the reduced 24nt-siRNA clusters provided in **Table S11** or the hypo CHH DMRs provided in **Table S12** showing the relationships between loci affected in the indicated single, double, and quadruple mutants relative to *pol-iv*. For readability, only overlaps >20 are labeled. A small number of overlaps are not shown due to spatial constraints, but unscaled Venn diagrams showing all the overlaps are present in **Fig. S5B**. For the 24nt-siRNA clusters, the Venn diagrams for the single and double mutants are also shown in **Fig. 3C**. (**B** and **D**) Boxplots showing 24nt-siRNA and DNA methylation levels at reduced 24nt-siRNA clusters (**B**) or showing DNA methylation levels at hypo CHH and CHG DMRs (**D**). Above each plot, the numbers (n) of reduced 24nt-siRNA clusters identified for each mutant are indicated and biological replicates for the WT controls are designated as WT1, WT2, and WT3, with the average signal from these replicates designated as the WT_av_.

**Figure S8. Relationships between 24nt-siRNA and DNA methylation levels for *clsy* mutants in ovule (Ov) tissue**. (**A** and **C**) Scaled Venn diagrams based on the reduced 24nt-siRNA clusters provided in **Table S11** or the hypo CHH DMRs provided in **Table S12** showing the relationships between loci affected in the indicated single, double, and quadruple mutants relative to *pol-iv*. For readability, only overlaps >20 are labeled. A small number of overlaps are not shown due to spatial constraints, but unscaled Venn diagrams showing all the overlaps are present in **Fig. S5B**. For the 24nt-siRNA clusters, the Venn diagrams for the single and double mutants are also shown in **Fig. 3C**. (**B** and **D**) Boxplots showing 24nt-siRNA and DNA methylation levels at reduced 24nt-siRNA clusters (**B**) or showing DNA methylation levels at hypo CHH and CHG DMRs (**D**). Above each plot, the numbers (n) of reduced 24nt-siRNA clusters identified for each mutant are indicated and biological replicates for the WT controls are designated as WT1, WT2, and WT3, with the average signal from these replicates designated as the WT_av_.

**Figure S9. Relationships between 24nt-siRNA and DNA methylation levels for *clsy* mutants in flower (Fl) tissue**. (**A**) Scaled Venn diagrams based on the reduced 24nt-siRNA clusters provided in **Table S11** showing the relationships between loci affected in the indicated single, double, and quadruple mutants relative to *pol-iv* mutants. For readability, only overlaps >20 are labeled. A small number of overlaps are not shown due to spatial constraints, but unscaled Venn diagrams showing all the overlaps are present in **Fig. S5B**. For the 24nt-siRNA clusters, the Venn diagrams for the single and double mutants are also shown in **Fig. 3C**. (**B**) Boxplots showing 24nt-siRNA and DNA methylation levels at reduced 24nt-siRNA clusters for the *clsy* mutants indicated above each set of plots. Above each plot, the numbers (n) of reduced 24nt-siRNA clusters identified for each mutant are indicated and biological replicates for the WT controls are designated as WT1, WT2, and WT3, with the average signal from these replicates designated as the WT_av_. Methylation levels for hypo CHH and CHG DMRs in the *clsy* and *pol-iv* mutants are published in Zhou *et al*.^34^.

**Figure S10. Assessment of DNA methylation levels at siren loci and motifs under CLSY3 peaks**. (**A**) Boxplots showing DNA methylation levels at siren 24nt-siRNA clusters (n=133) in the indicated genotypes and tissues. (**B**) Top four motifs identified from the CLSY3 ChIP, including their p-values and the percent of each motif found in target regions (CLSY3 peaks n=102) or background regions (TAIR10 genome n=43,700). (**C**) Screenshots showing the levels of DNA methylation (% methylation in each context), 24nt-siRNAs, and CLSY3 or Pol-IV enrichment from flower tissue in the indicated genotypes at representative CLSY3 peaks. Track scales are the same for all three sites and are indicated in brackets. The regions corresponding to different features are indicated below. For the Pol-IV ChIP data the IP subscripts indicate the genetic background (*e*.*g*. IP_c3_ indicates the *clsy3* background).

**Figure S11. Chromosomal Distributions of 24nt-siRNA clusters**. (**A**) Distributions of reduced 24nt-siRNA clusters, in the indicated genotypes and tissues, along the 5 chromosomes. The pericentromeric heterochromatin, as designated in Yelina *et al*.^66^, is marked in red and denoted by vertical dashed lines. Scale bars for x-axis (Mb) and y-axis (clusters/100kb bin) are indicated in the lower left corner of each set. Note that ovules are at 2x relative to the other tissues. Data for the *clsy1,2* and *clsy3,4* clusters are also shown in **Fig. 5E**.

**Figure S12. 24nt-siRNA and DNA methylation levels in relation to the 10 24nt-siRNA classes**. (**A**) Boxplots showing 24nt-siRNA and DNA methylation levels in all sequence contexts (% all mC) at reduced 24nt-siRNA clusters grouped by the ten classes defined in **Fig. 2C** (upper) or at hypo CHH DMRs within these clusters (lower) for the genotypes and tissues indicated below. The numbers (n) of clusters or DMRs are indicated and biological replicates for the WT controls are designated as WT1, WT2, and WT3, with the average signal from these replicates designated as the WT_av_.

## Supplementary Tables

Table S1. Summary of mRNA-seq data

Table S2. Summary of smRNA-seq data

Table S3. Summary of MethylC-seq data

Table S4. 21-24nt small RNA clusters

Table S5. 24nt-siRNA cluster lists and summary

Table S6. 24nt-siRNA heatmap clustering and summary

Table S7. DMRs between WT tissues and summary

Table S8. DNA methylation heatmap clustering and summary

Table S9. *clsy* upregulated DEGs overlapping with *pol-iv* and summary

Table S10. *pol-iv* DE analysis and summary

Table S11. DE 24nt-siRNA clusters and summary

Table S12. hypo DMRs and summary

Table S13. Summary of ChIP-seq data

Table S14. ChIP analysis and summary Table S15. Primers table

## Acknowledgements

We thank colleagues and lab members for helpful comments and discussions. This work was supported by the NIH (GM112966) and Hearst Foundation to J. Law. Dr. Ming Zhou was supported by the 111 Project (Grant B14027), the Fundamental Research Funds for the Central Universities at Zhejiang University (2020QNA6006), the Hundred-Talent Program of Zhejiang University, a Pioneer Fund Postdoctoral Award, and a postdoctoral fellowship from the Paul F. Glenn Center for Biology of Aging Research at the Salk Institute. This work was also supported by the NGS Core Facility and the Integrative Genomics and Bioinformatics Core Facility at the Salk Institute with funding from NIH-NCI CCSG: P30 014195, the Paul F. Glenn Center for Biology of Aging Research at the Salk Institute, the Chapman Foundation and the Helmsley Charitable Trust.

## Notes

### Competing Interest Statement

The authors have declared no competing interest.

### Summary of Updates

The six main figures were replaced with versions that start lower on the page to avoid overlaps with the bioRxiv generated header.

