## Supplemental Figures for "The CLASSY family controls tissue-specific DNA methylation patterns in Arabidopsis"

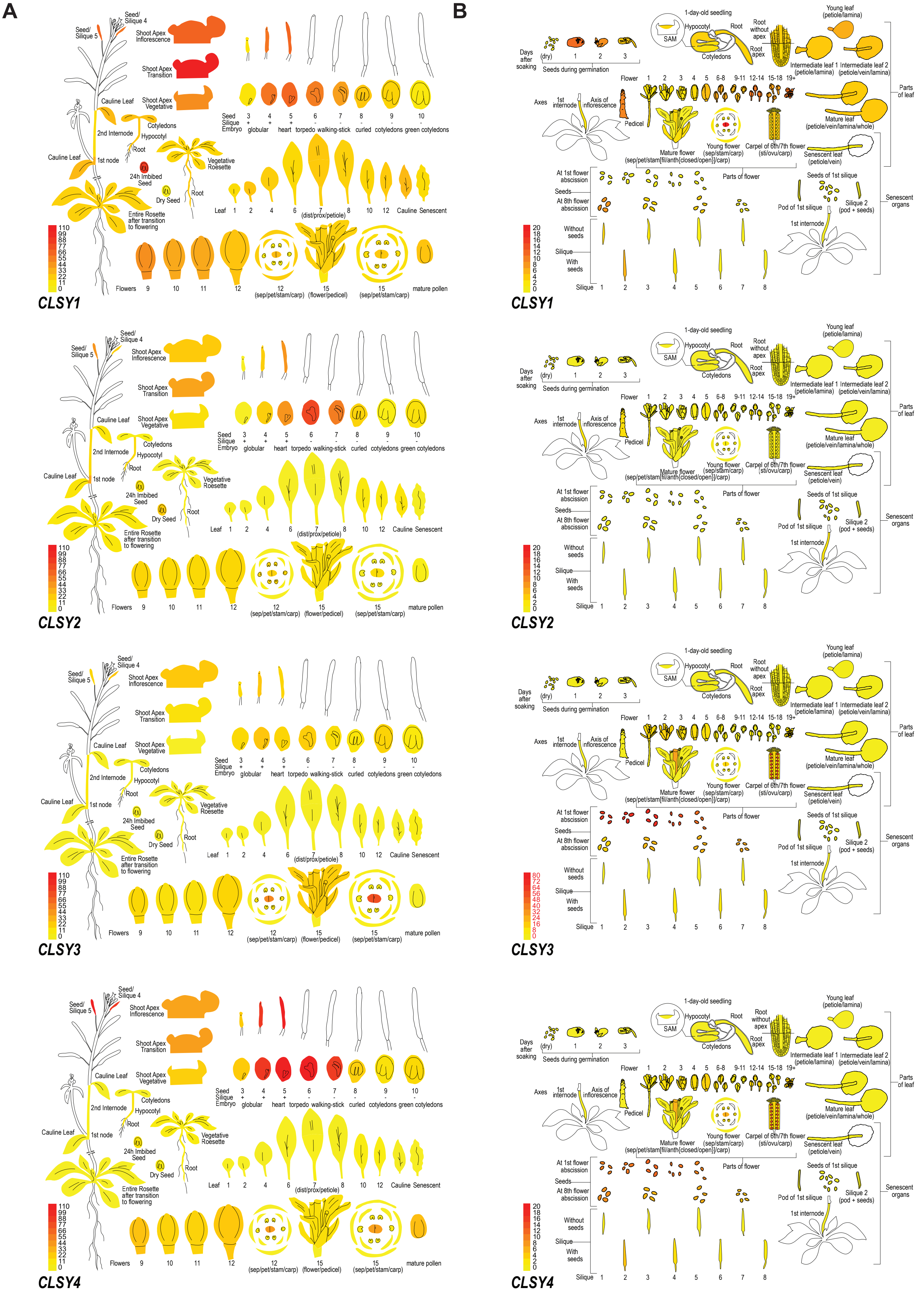


**Figure S1. Visualization of *CLSY* expression patterns using ePlant**[35]**.** Relative expression levels of the four *CLSY* genes from two Plant eFP viewers: (**A**) AtGenExpress[69,70] and (**B**) RNA-seq[68]. For comparison, the color scales are equal for panels **A** and **B** with the exception of *CLSY3*, which is on a higher scale, as indicated by the red text on the color bar. All tissues and organs are labeled as in the ePlant viewers[35]. Images are not to scale.

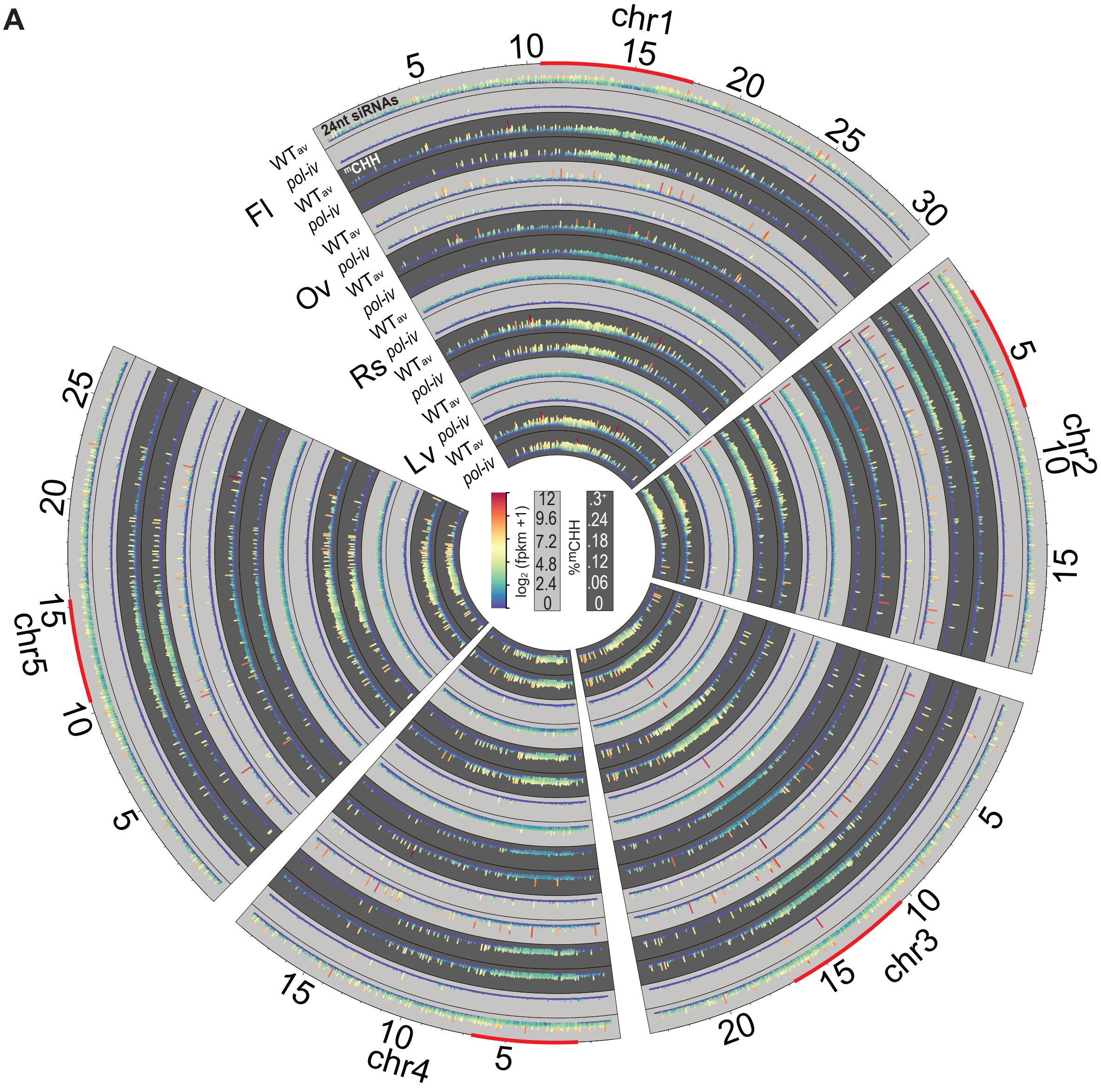


**Figure S2. Global characterization of 24nt-siRNA and CHH methylation patterns across tissues.** (**A**) Circular genome view showing the patterns of 24nt-siRNAs (light grey background) and CHH methylation (dark grey background) across all five chromosomes (chr1-5) in 5kb bins based on the $WT_{av}$ expression levels from each tissue. The color scales for the data sets are as indicated in the center of the plot and the tracks are labeled every 5Mbs, with the pericentromeric heterochromatin, as designated in Yelina *et al.*[66], marked in red along the outer circle. For CHH methylation, between 1 and 3 bins, depending on the tissue, had values over 0.3 ($0.3^{+}$), but were capped at this value to facilitate visualization on a genomic scale.

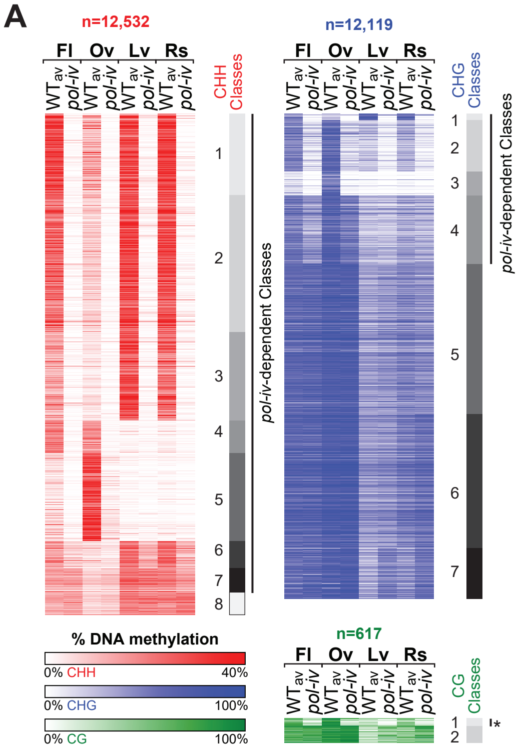


**Figure S3. Tissue specific comparison of DNA methylation patterns.** (**A**) Scaled heatmaps showing the designation of 8 CHH, 7 CHG, and 2 GC classes of hypo DMRs, respectively, based on the methylation levels ($WT_{av}$ and *pol-iv*) for each tissue at the master set of WTvsWT DMRs for each context (**Table S7**). Classes showing clear reductions in methylation in the *pol-iv* mutant are indicated as *pol-iv*-dependent. For CG DMRs, this is indicated by an asterisk.

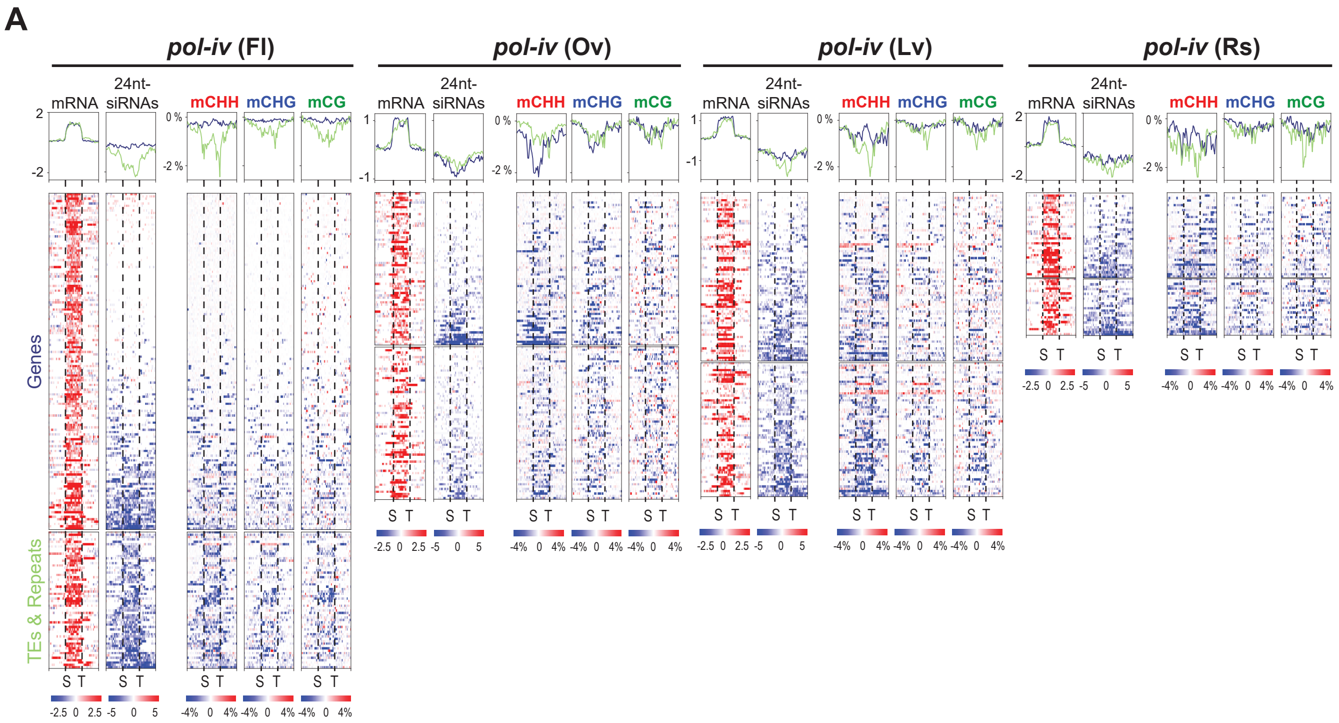


**Figure S4. Assessment of 24nt-siRNA and DNA methylation levels surrounding differentially expressed transcripts.** (**A**) Heatmaps and profile plots showing the expression of all *pol-iv* upregulated transcripts ($log_2$ FC ≥ 1 and FDR ≤ 0.05; as shown in (**Fig. 2G**)) as well as the corresponding 24nt-siRNA and DNA methylation levels at these same loci. For the mRNA and 24nt-siRNA analyses, the $log_2$ fold changes in expression in *pol-iv* mutants relative to wild-type controls are plotted and for the DNA methylation analysis, the difference in the percent methylation between *pol-iv* mutants and wild-type controls is plotted. Color bars indicating the scales are shown below. The heatmaps include 2kb flanking the transcription start site (S) and the transcription termination site (T) and were ranked based on the 24nt-siRNA and mCHH values for each tissue. The profiles for the genes or TE/repeats are shown in blue and light green, respectively, above each heatmap. For flower tissue, the data from Zhou *et al*.[34] was reanalyzed using an updated genome annotation (**Source Data 1**).

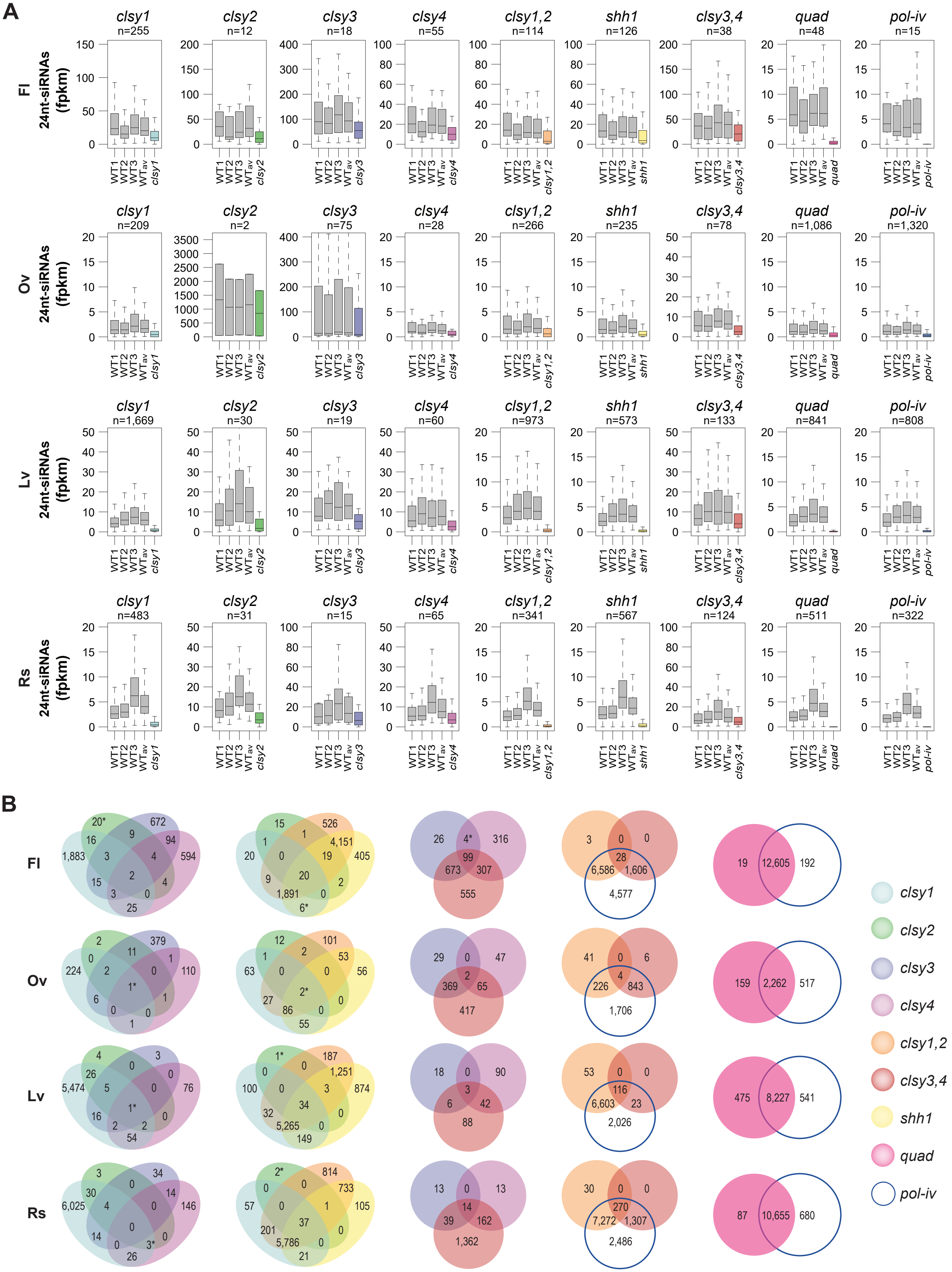


**Figure S5. Additional analysis of 24nt-siRNA clusters affected in the different tissues and mutant backgrounds.** (**A**) Boxplots showing 24nt-siRNA levels at clusters for each tissue and genotype that overlap with hypo CHH DMR(s) and are reduced (>25%) compared to tissue matched WT controls, but were not initially detected as DE clusters due to the FC and p-value thresholds. Here, and for all subsequent boxplots, the graphs show the interquartile range (IQR), with the median shown as a black line and the whiskers corresponding to 1.5 times the IQR. Above each plot, the numbers (n) of clusters are indicated and biological replicates for the WT controls are designated as WT1, WT2, and WT3, with the average signal from these replicates designated as the $WT_{av}$. (**B**) Unscaled Venn diagrams showing the overlaps between 24nt-siRNA clusters affected in each mutant combination and tissue type. Mutants are colored as indicated on the right. * indicates overlaps not included in the scaled Venn diagrams due to spatial constraints.

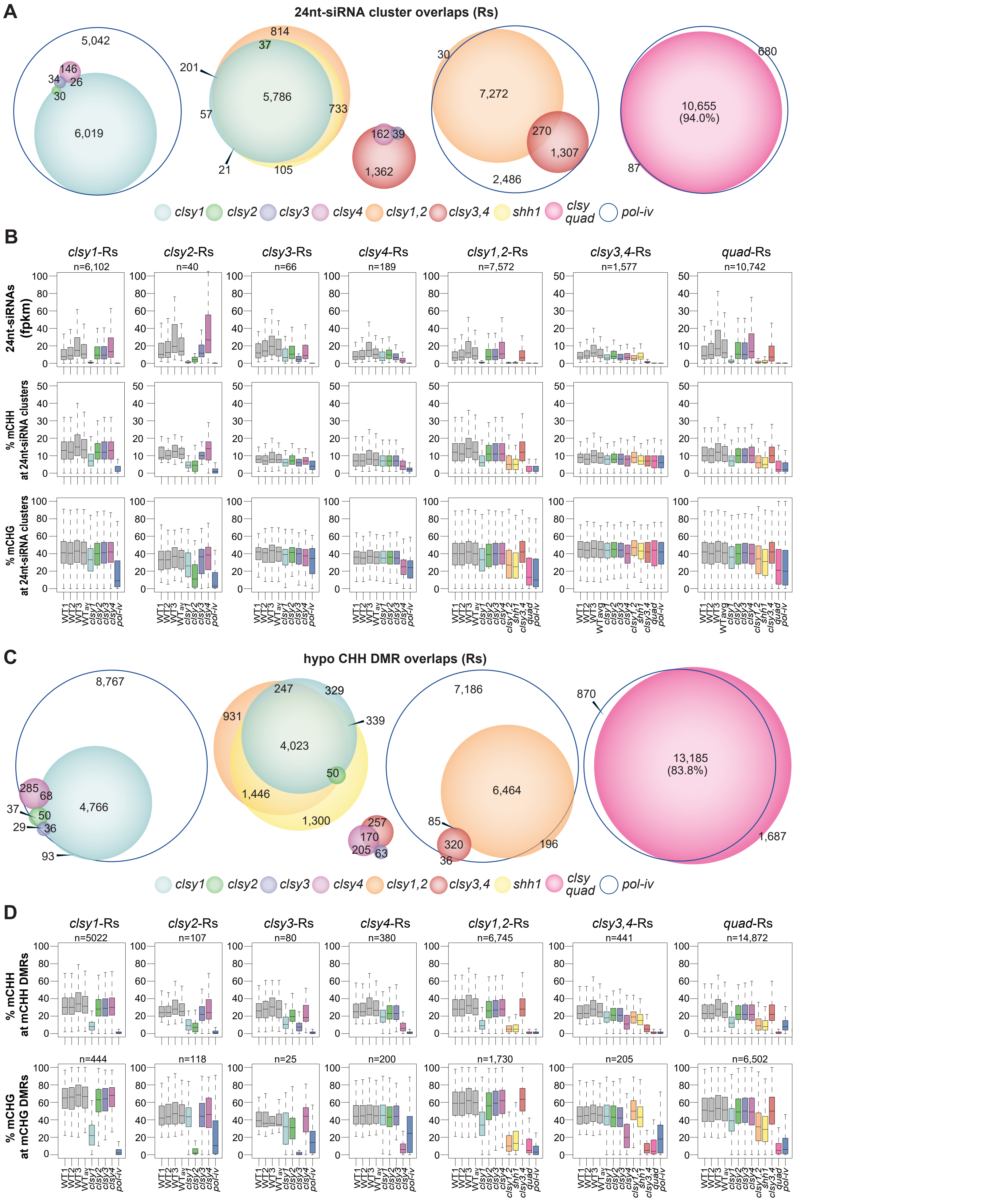


**Figure S6. Relationships between 24nt-siRNA and DNA methylation levels for *clsy* mutants in rosette (Rs) tissue.** (**A** and **C**) Scaled Venn diagrams based on the reduced 24nt-siRNA clusters provided in **Table S11** or the hypo CHH DMRs provided in **Table S12** showing the relationships between loci affected in the indicated single, double, and quadruple mutants relative to *pol-iv*. For readability, only overlaps >20 are labeled. A small number of overlaps are not shown due to spatial constraints, but unscaled Venn diagrams showing all the overlaps are present in **Fig. S5B**. For the 24nt-siRNA clusters, the Venn diagrams for the single and double mutants are also shown in **Fig. 3C**. (**B** and **D**) Boxplots showing 24nt-siRNA and DNA methylation levels at reduced 24nt-siRNA clusters (**B**) or showing DNA methylation levels at hypo CHH and CHG DMRs (**D**). Above each plot, the numbers (n) of reduced 24nt-siRNA clusters identified for each mutant are indicated and biological replicates for the WT controls are designated as WT1, WT2, and WT3, with the average signal from these replicates designated as the $WT_{av}$.

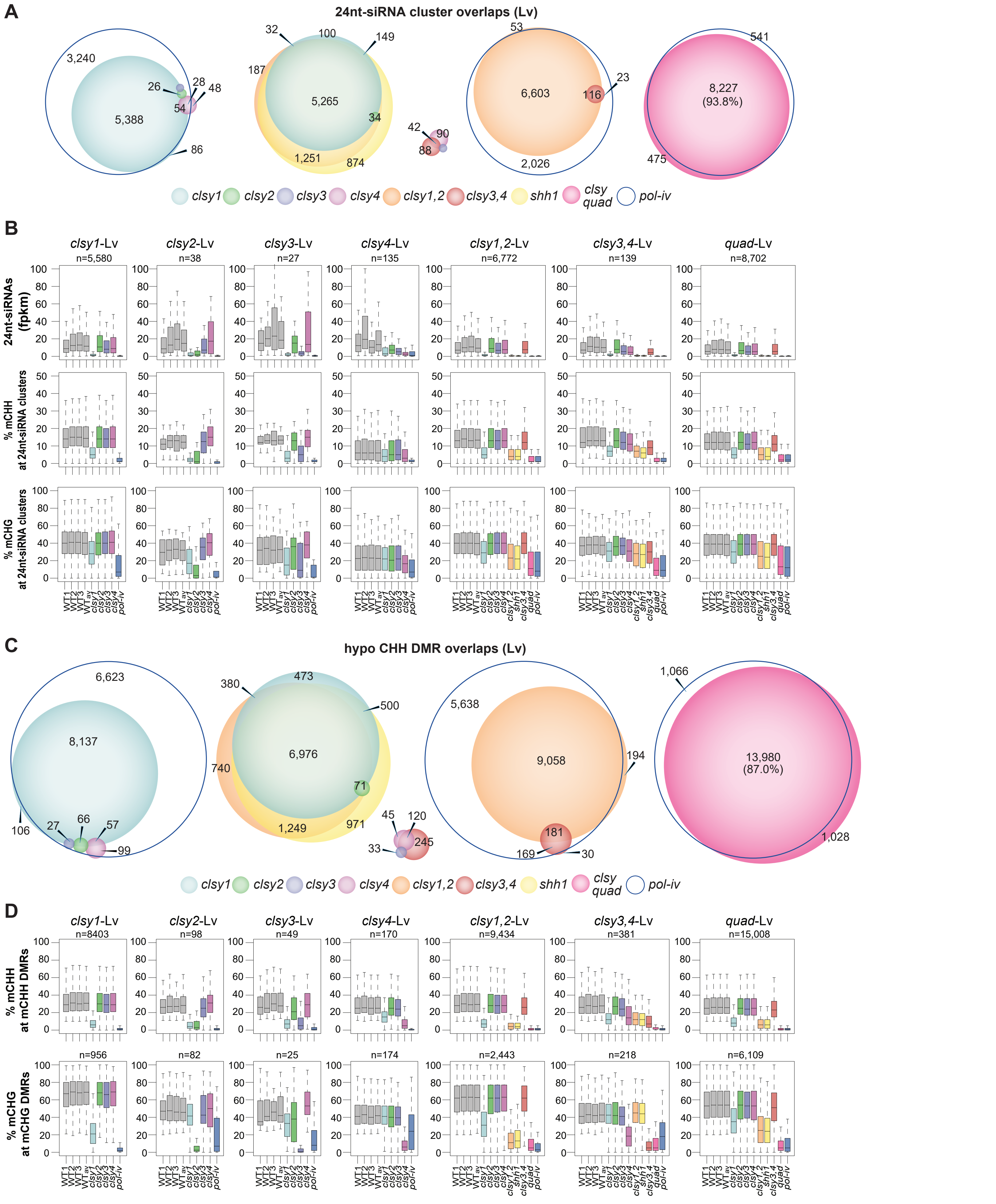


**Figure S7. Relationships between 24nt-siRNA and DNA methylation levels for *clsy* mutants in leaf (Lv) tissue.** (**A** and **C**) Scaled Venn diagrams based on the reduced 24nt-siRNA clusters provided in **Table S11** or the hypo CHH DMRs provided in **Table S12** showing the relationships between loci affected in the indicated single, double, and quadruple mutants relative to *pol-iv*. For readability, only overlaps >20 are labeled. A small number of overlaps are not shown due to spatial constraints, but unscaled Venn diagrams showing all the overlaps are present in **Fig. S5B**. For the 24nt-siRNA clusters, the Venn diagrams for the single and double mutants are also shown in **Fig. 3C**. (**B** and **D**) Boxplots showing 24nt-siRNA and DNA methylation levels at reduced 24nt-siRNA clusters (**B**) or showing DNA methylation levels at hypo CHH and CHG DMRs (**D**). Above each plot, the numbers (n) of reduced 24nt-siRNA clusters identified for each mutant are indicated and biological replicates for the WT controls are designated as WT1, WT2, and WT3, with the average signal from these replicates designated as the $WT_{av}$.

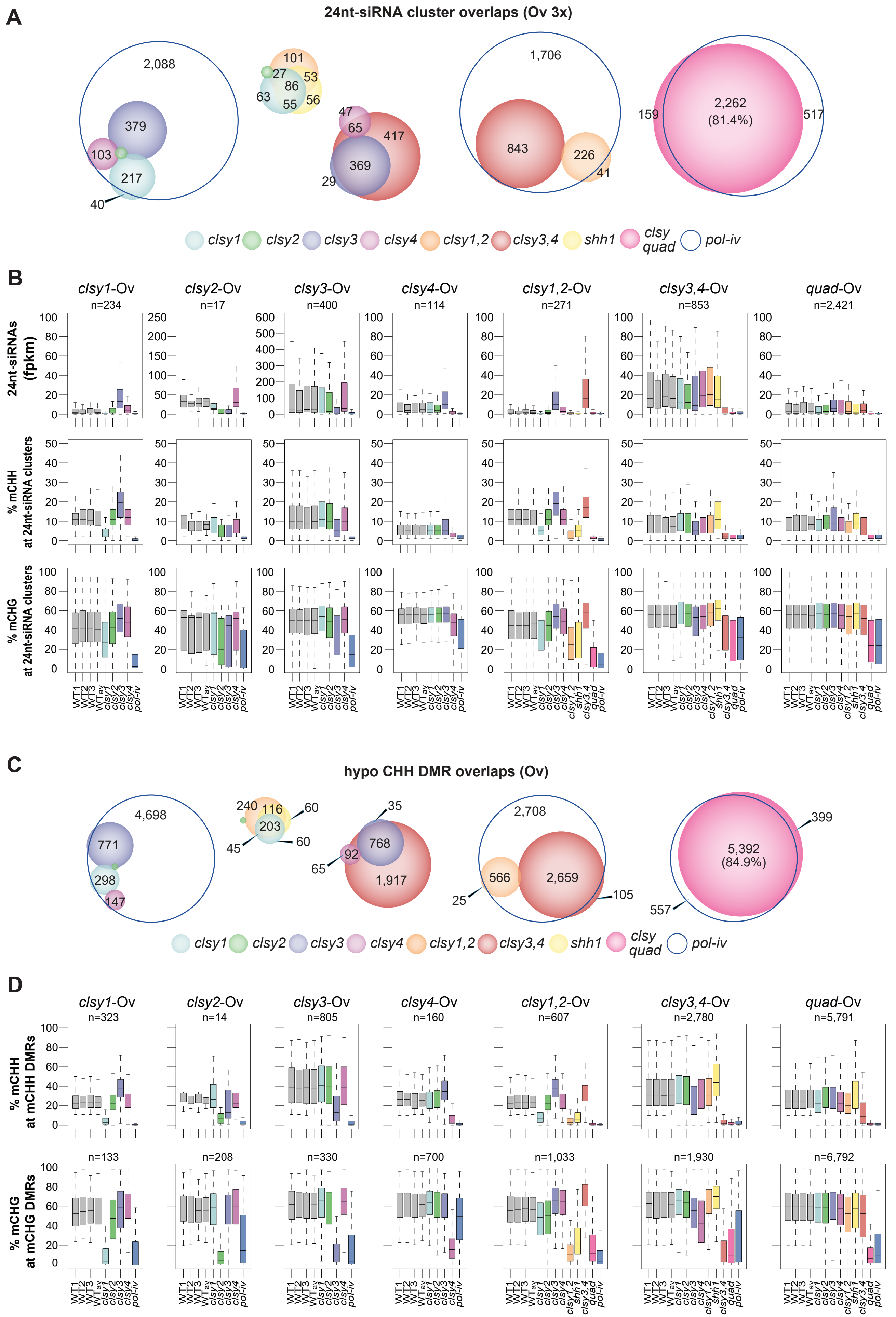


**Figure S8. Relationships between 24nt-siRNA and DNA methylation levels for *clsy* mutants in ovule (Ov) tissue.** (**A** and **C**) Scaled Venn diagrams based on the reduced 24nt-siRNA clusters provided in **Table S11** or the hypo CHH DMRs provided in **Table S12** showing the relationships between loci affected in the indicated single, double, and quadruple mutants relative to *pol-iv*. For readability, only overlaps >20 are labeled. A small number of overlaps are not shown due to spatial constraints, but unscaled Venn diagrams showing all the overlaps are present in **Fig. S5B**. For the 24nt-siRNA clusters, the Venn diagrams for the single and double mutants are also shown in **Fig. 3C.** (**B** and **D**) Boxplots showing 24nt-siRNA and DNA methylation levels at reduced 24nt-siRNA clusters (**B**) or showing DNA methylation levels at hypo CHH and CHG DMRs (**D**). Above each plot, the numbers (n) of reduced 24nt-siRNA clusters identified for each mutant are indicated and biological replicates for the WT controls are designated as WT1, WT2, and WT3, with the average signal from these replicates designated as the $WT_{av}$.

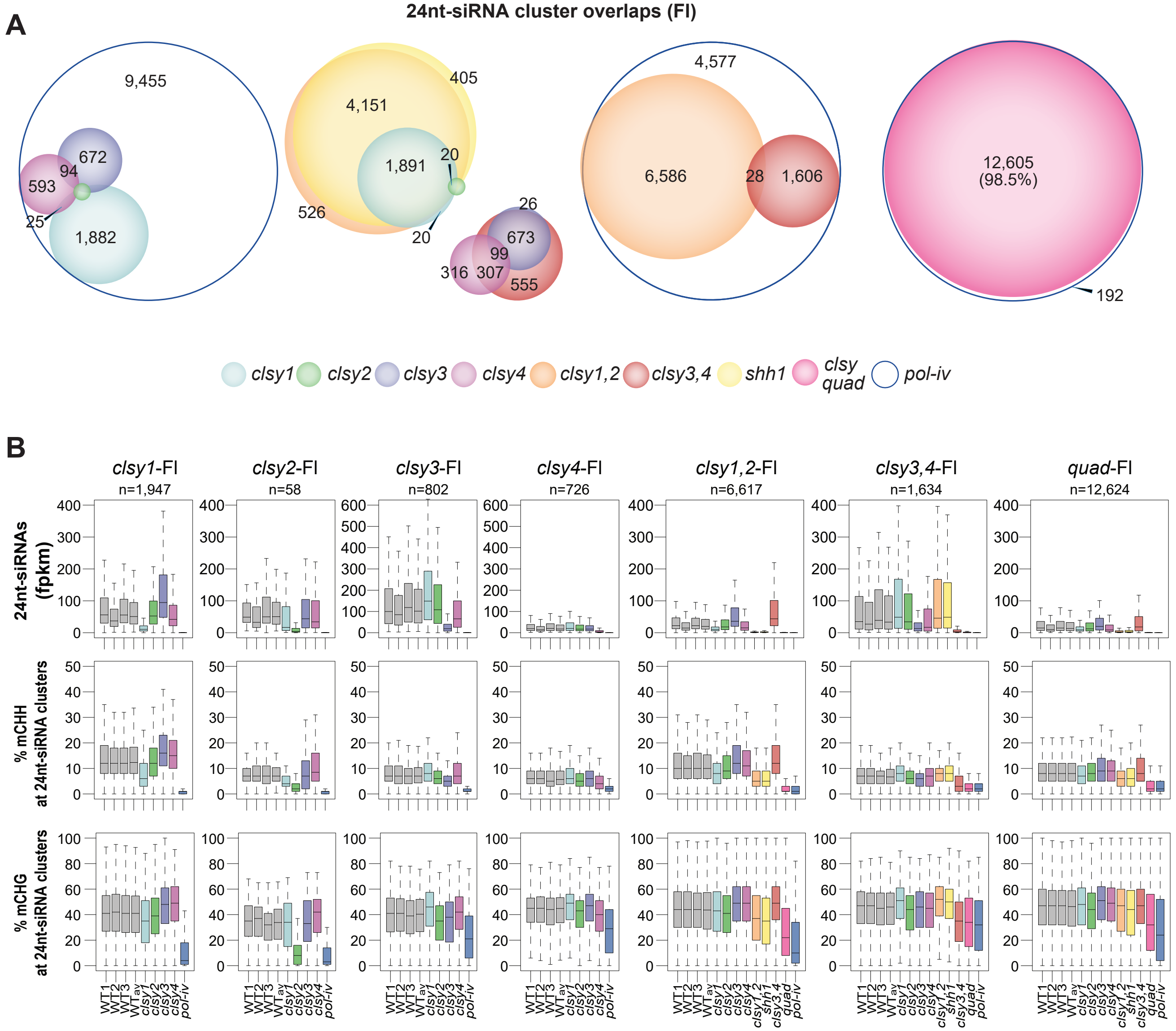


**Figure S9. Relationships between 24nt-siRNA and DNA methylation levels for *clsy* mutants in flower (Fl) tissue.** (**A**) Scaled Venn diagrams based on the reduced 24nt-siRNA clusters provided in **Table S11** showing the relationships between loci affected in the indicated single, double, and quadruple mutants relative to *pol-iv* mutants. For readability, only overlaps >20 are labeled. A small number of overlaps are not shown due to spatial constraints, but unscaled Venn diagrams showing all the overlaps are present in **Fig. S5B**. For the 24nt-siRNA clusters, the Venn diagrams for the single and double mutants are also shown in **Fig. 3C**. (**B**) Boxplots showing 24nt-siRNA and DNA methylation levels at reduced 24nt-siRNA clusters for the *clsy* mutants indicated above each set of plots. Above each plot, the numbers (n) of reduced 24nt-siRNA clusters identified for each mutant are indicated and biological replicates for the WT controls are designated as WT1, WT2, and WT3, with the average signal from these replicates designated as the $WT_{av}$. Methylation levels for hypo CHH and CHG DMRs in the *clsy* and *pol-iv* mutants are published in Zhou *et al*.[34].

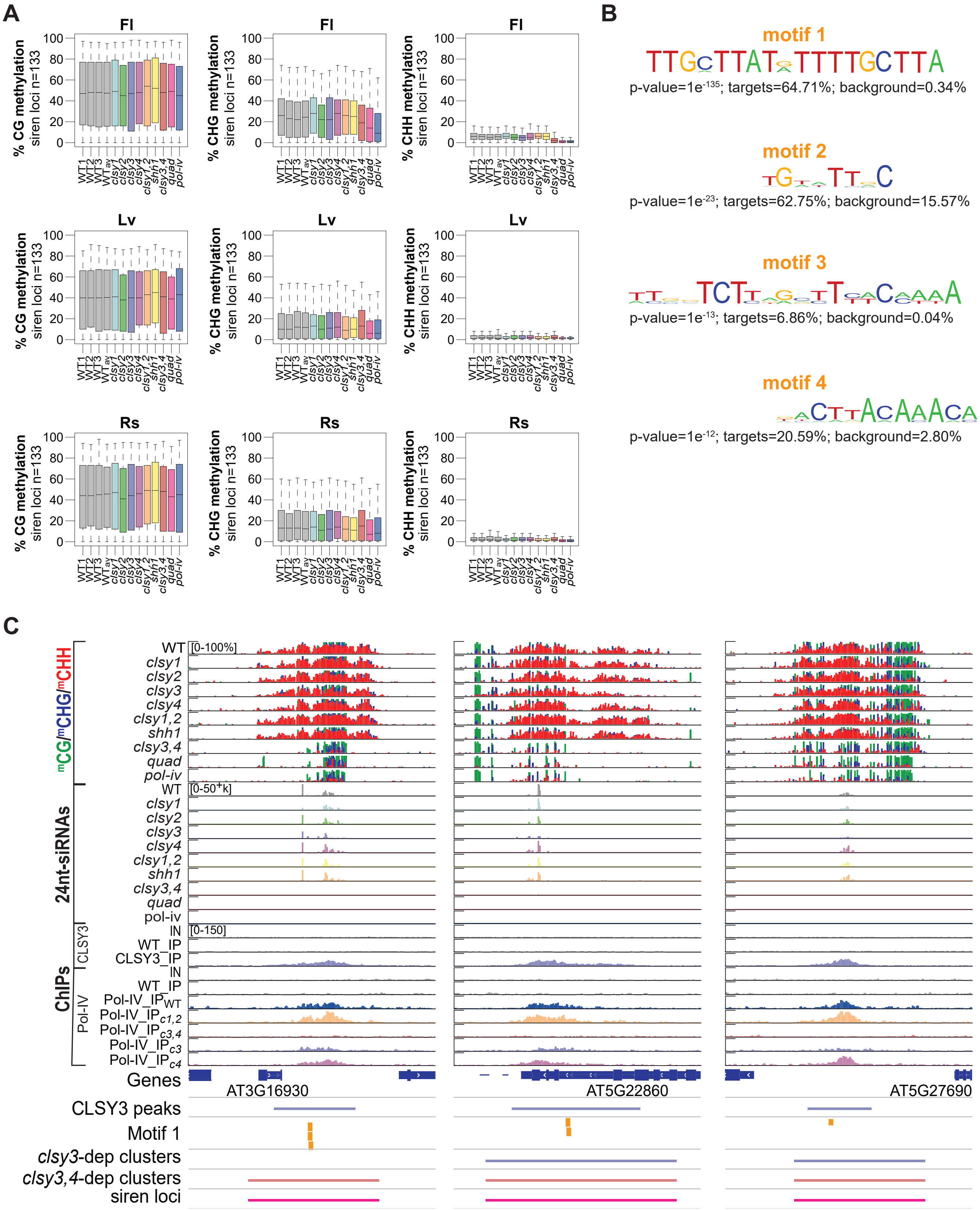

**Figure S10. Assessment of DNA methylation levels at siren loci and motifs under CLSY3 peaks.** (**A**) Boxplots showing DNA methylation levels at siren 24nt-siRNA clusters (n=133) in the indicated genotypes and tissues. (**B**) Top four motifs identified from the CLSY3 ChIP, including their p-values and the percent of each motif found in target regions (CLSY3 peaks n=102) or background regions (TAIR10 genome n=43,700). (**C**) Screenshots showing the levels of DNA methylation (% methylation in each context), 24nt-siRNAs, and CLSY3 or Pol-IV enrichment from flower tissue in the indicated genotypes at representative CLSY3 peaks. Track scales are the same for all three sites and are indicated in brackets. The regions corresponding to different features are indicated below. For the Pol-IV ChIP data the IP subscripts indicate the genetic background (e.g. $IP_{c3}$ indicates the *clsy3* background).

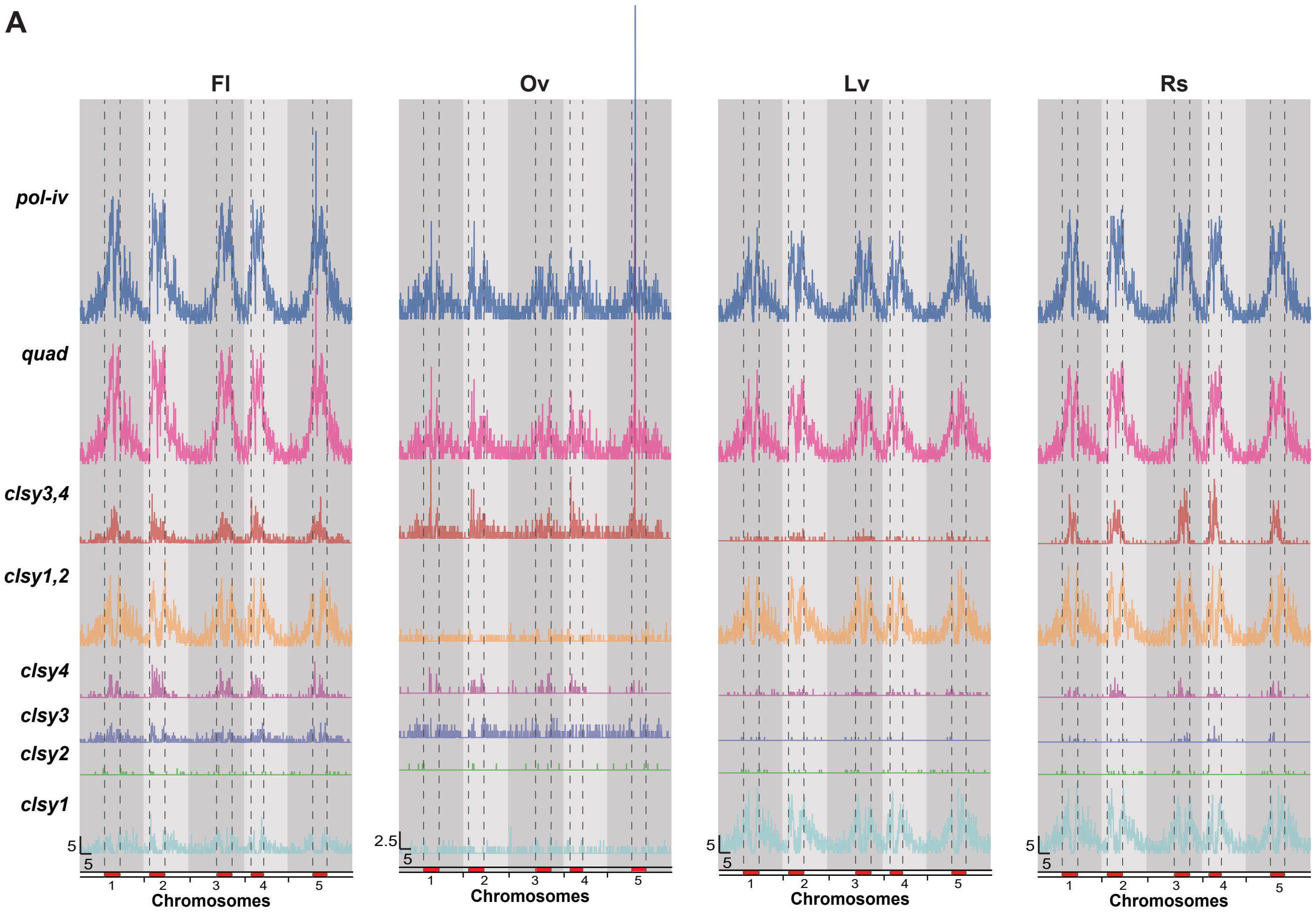


**Figure S11. Chromosomal Distributions of 24nt-siRNA clusters.** (**A**) Distributions of reduced 24nt-siRNA clusters, in the indicated genotypes and tissues, along the 5 chromosomes. The pericentromeric heterochromatin, as designated in Yelina *et al.*[66], is marked in red and denoted by vertical dashed lines. Scale bars for x-axis (Mb) and y-axis (clusters/100kb bin) are indicated in the lower left corner of each set. Note that ovules are at 2x relative to the other tissues. Data for the *clsy1,2* and *clsy3,4* clusters are also shown in **Fig. 5E**.

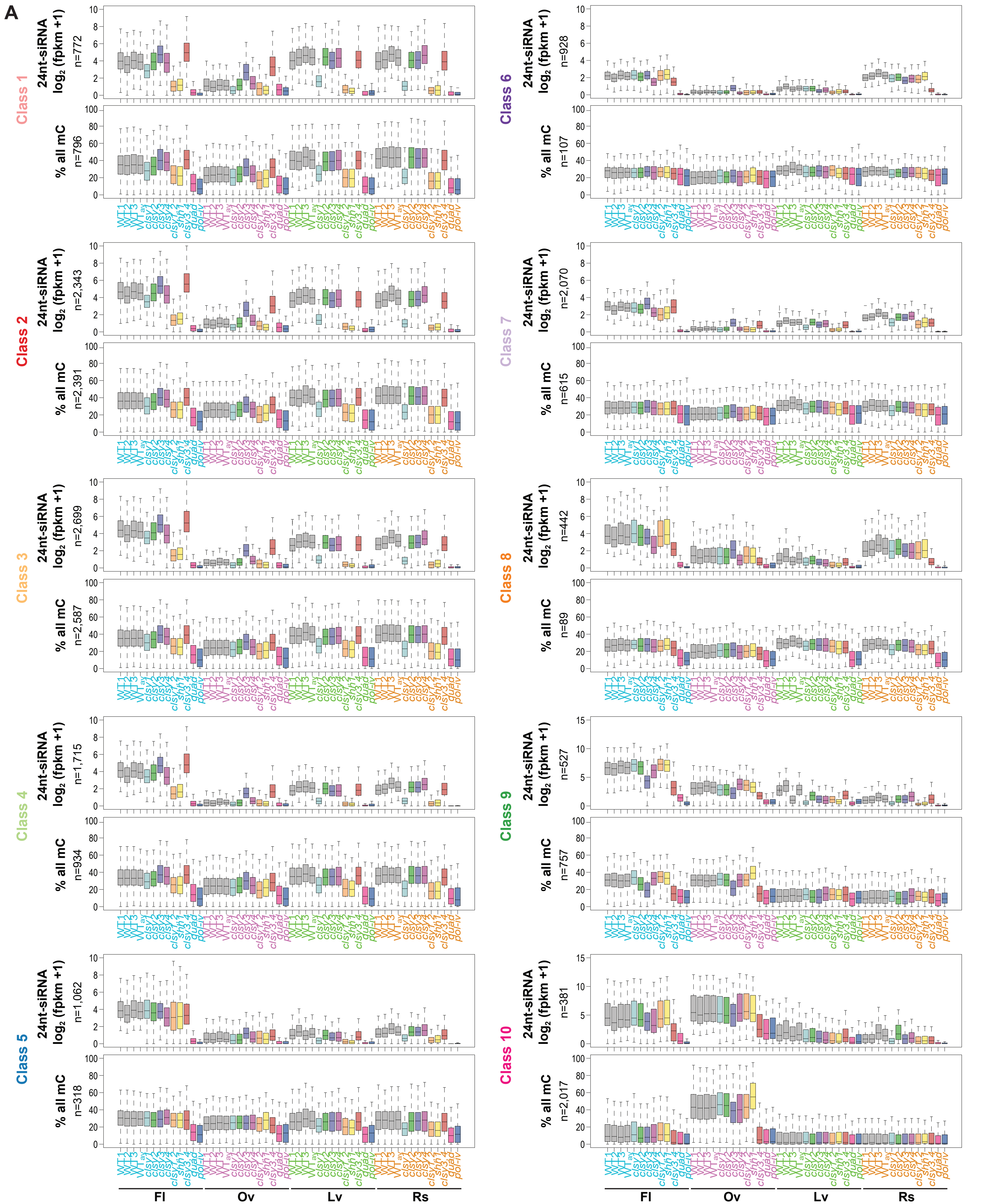


**Figure S12. 24nt-siRNA and DNA methylation levels in relation to the 10 24nt-siRNA classes.** (**A**) Boxplots showing 24nt-siRNA and DNA methylation levels in all sequence contexts (% all mC) at reduced 24nt-siRNA clusters grouped by the ten classes defined in **Fig. 2C** (upper) or at hypo CHH DMRs within these clusters (lower) for the genotypes and tissues indicated below. The numbers (n) of clusters or DMRs are indicated and biological replicates for the WT controls are designated as WT1, WT2, and WT3, with the average signal from these replicates designated as the $WT_{av}$.
